# Drought tolerance as an evolutionary precursor to frost and winter tolerance in grasses

**DOI:** 10.1101/2024.06.29.601311

**Authors:** Laura Schat, Marian Schubert, Siri Fjellheim, Aelys M. Humphreys

**Affiliations:** Department of Ecology, Environment and Plant Sciences, Stockholm University, Stockholm, Sweden; Bolin Centre for Climate Research, Stockholm University, Stockholm, Sweden; Department of Plant Sciences, Norwegian University of Life Sciences, Ås, Norway

**Keywords:** biome transition, correlated evolution, GBIF data, Köppen-Geiger, rate-heterogeneity, severe winter

## Abstract

Accumulating evidence is suggesting more frequent tropical-to-temperate transitions than previously thought. This raises the possibility that biome transitions could be facilitated by precursor traits. A wealth of ecological, genetic and physiological evidence suggests overlap between drought and frost stress responses, but the origin of this overlap, i.e. the evolution of these responses relative to each other, is poorly known. Here, we test whether adaptation to frost and/or severe winters in grasses (Poaceae) was facilitated by ancestral adaptation to drought. We used occurrence patterns across Köppen-Geiger climate zones to classify species as drought, frost and/or winter tolerant, followed by comparative analyses. Ancestral state reconstructions revealed different evolutionary trajectories in different clades, suggesting both drought-first and frost-first scenarios. A model of correlated evolution was not supported when transition rate heterogeneity was taken into account or compared to traits simulated under independent evolution. Our findings provide some support for ancestral drought tolerance facilitating transitions to cold, temperate biomes, at least in some clades. Different scenarios in different clades is consistent with present-day grasses being either cold or drought specialists, possibly as a consequence of trade-offs between different stress tolerance responses.

## Introduction

A general pattern of plant diversity is that it is highest in the warm, wet tropics and declines towards the poles, where conditions become colder and more seasonally variable. Most major angiosperm clades originated in the tropics, as tropical climates predate temperate ones (e.g. Judd et al. 1994; Ricklefs and Renner 1994; Zanne et al. 2014). When temperate biomes expanded after the Eocene, tropical lineages in emerging temperate biomes had to adapt to drier, colder and more seasonal climates to persist (Eldrett et al., 2009; Kerkhoff et al., 2014; Zachos et al., 2001). Yet, many clades do not occur at higher latitudes, despite plenty of evolutionary opportunity and time to become established there (Donoghue, 2008). Causes of this latitudinal diversity gradient have been the subject of a debate spanning decades (Fine & Ree, 2006; Hillebrand, 2004; Jansson et al., 2013; Kerkhoff et al., 2014; Mittelbach et al., 2007; Rohde, 1992; Wiens & Donoghue, 2004; Wiens & Graham, 2005).

One prevalent explanation assumes that tropical-to-temperate transitions have been infrequent because adapting to novel, and especially freezing, climates is difficult (e.g. Wiens and Donoghue 2004; Wiens and Graham 2005; Donoghue 2008; Wiens et al. 2010; Körner 2016). As a result, temperate clades are expected to be clustered phylogenetically and nested within tropical clades (Kerkhoff et al., 2014).

Previous reviews have suggested that approximately 40% -50% of all angiosperm families have temperate lineages (Preston & Sandve, 2013; Stevens, 2001), and indeed, on a global scale, biome shifts are thought to have been rare (Crisp et al., 2009). However, recent research is indicating that tropical-to-temperate transitions may have been more frequent than currently thought (Donoghue & Edwards, 2014; Jansson et al., 2013; Nürk et al., 2018; Zizka et al., 2020), apparently having occurred multiple times independently, even within single families (e.g. Poaceae; Edwards and Smith 2010, Schubert et al. 2020; Amaranthaceae; Cousins-Westerberg et al. 2023) and genera (e.g. *Viburnum*; Schmerler et al. 2012; Spriggs et al. 2015). This raises the possibility that such transitions are not necessarily as difficult as widely assumed, which in turn could imply that biome transitions are facilitated by evolutionary precursor trait(s) (also referred to as “pre-adaptations” or “enabler traits”; Edwards and Donoghue 2013; Donoghue and Edwards 2014). For tropical-to-temperate transitions, such precursors could be related to ancient stress tolerance mechanisms that have been repurposed for the specific challenges of a cold, temperate climate (Preston & Sandve, 2013). If so, we would expect adaptations to frost and long winters to have arisen more frequently in clades with such evolutionary precursor(s) than in clades without them, leading to a pattern of multiple apparent “independent” transitions to a temperate climate in those clades.

Temperate plants are confronted by a range of abiotic stresses. For example, frost impacts plants on many levels, ranging from dehydration stress caused by the lack of fluid water, to damage to the structural integrity of cells and biomembranes caused by ice crystal formation (Preston & Sandve, 2013; Sakai & Larcher, 1987). Additionally, plant growth slows down at temperatures below 5 °C and comes to a near standstill at freezing (Körner, 2016; Nievola et al., 2017). Overcoming these abiotic stresses and evolving frost tolerance requires major physiological innovations (Donoghue, 2008; Lancaster & Humphreys, 2020; Sakai & Larcher, 1987; Wiens & Donoghue, 2004).

Plants in freezing areas can be exposed to frost episodically (e.g. diurnally), or periodically (i.e. during prolonged periods of seasonal or even permanent frost). Temperatures can drop below freezing occasionally in the tropics (e.g. on tropical mountains) and across the temperate zone, but plants in areas with long, cold winters are in addition confronted by seasonal variation in precipitation, temperature and day length and must endure long periods of low (but not necessarily freezing) temperatures. Frost tolerance alone is thus insufficient to survive cold, temperate winters, which requires a suite of physiological and phenological adjustments in response to seasonal change (cold acclimation and vernalisation) and a short (and possibly cold) growing season. To capture some of the stress tolerance mechanisms required by temperate plants, we separately consider exposure to both frost and severe winters in this study.

There is strong ecological, genetic and physiological evidence that adaptations to low temperature and drought stress are not independent. On a cellular level, the dehydration caused by extracellular ice is virtually identical to dehydration stress caused by drought (Körner, 2016; Sakai & Larcher, 1987), and several genes involved in responses to cold and frost stress are also expressed during drought stress (Birkeland et al., 2021; Das et al., 2023; Schubert, Grønvold, et al., 2019; Zhong et al., 2018). Both periodic drought and winter present long periods without possibilities for photosynthesis, and plants need storage carbohydrates that can be hydrolysed to provide fuel to survive both of these conditions.

Further, plants from dry mountains are more frost tolerant than those from wet ones (Sierra-Almeida et al., 2016), and water-deficiency pre-treatment can increase frost tolerance whereas pre-treatment with heat does not affect subsequent frost tolerance (Sumner et al., 2022). However, while there is plenty of evidence for a correlation between responses to frost and drought stress, much less is known about how the overlap among different abiotic stress responses evolved. One hypothesis is that ancient stress tolerance mechanisms might have been beneficial to surviving in both cold and dry areas (Folk et al., 2020; Preston & Sandve, 2013). Another hypothesis is that lineages that transitioned to freezing areas might have been predisposed to do so thanks to precursor traits that are beneficial in a freezing environment but evolved in response to aridity in the tropics (e.g. small conduits and a herbaceous habit; Zanne et al. 2014).

In general, some strategy for avoiding dehydration is thought to be more ancient than any adaptation to low temperature stress. Early land plants (Embryophyta) transitioning to terrestrial environments ca. 500 million years (Ma) ago were most likely desiccation tolerant; thus desiccation tolerance is thought to be an ancestral trait in land plants and a key component of adaptation to life on land (Bowles et al., 2021; Oliver, 1917; Preston & Sandve, 2013; Sakai & Larcher, 1987). However, desiccation tolerance is less common in vascular plants (Tracheophyta); their origin saw the evolution of vascular tissue and other regulatory and morphological complexities that enabled them to tolerate dry conditions (Bowles et al., 2021; Harrison, 2017). Thus, desiccation tolerance was probably lost in the ancestor of vascular plants and replaced by early forms of drought tolerance (Bowles et al., 2021). In contrast, while cool-climate pockets may have been present in mid-latitude mountain areas in the Northern Hemisphere in the Eocene (56–34 Ma; Hagen et al. 2019), the emergence of cold and freezing environments of today is not thought to have begun until the late Eocene or early Oligocene (i.e. mainly in the past ca. 34 Ma; Eldrett et al., 2009; Liu et al., 2009; Pound & Salzmann, 2017; Zachos et al., 2001). Thus, strategies for dealing with low temperature stress across living land plants are thought to have evolved by independent repurposing of ancestral stress pathways (e.g. related to dehydration; Preston & Sandve, 2013; Schubert et al., 2020; Schubert, Grønvold, et al., 2019), rather than representing ancestral traits.

Here we explore the idea of existing stress responses facilitating adaptation to freezing environments in grasses (Poaceae), by testing whether there is evidence that drought tolerance could have acted as the evolutionary precursor that facilitated persistence in the emerging cold environments of the Oligocene. Poaceae is one of the five largest angiosperm families, with over 11,000 species and 800 genera distributed largely in two main clades: the BOP clade (Bambusoideae, Oryzoideae, Pooideae) and PACMAD clade (Panicoideae, Aristidoideae, Chloridoideae, Micrairoideae, Arundinoideae, Danthonioideae; Kellogg 2001; Clayton et al. 2006; Hodkinson 2018; Figure 1). They are well suited for testing our hypothesis of precursor traits due to their cosmopolitan distribution, occurring from moist tropical forests and tropical savannahs to arctic tundras and alpine meadows, including some of the harshest environments on Earth (e.g. hot and cold deserts, saline environments and Antarctica; Bannister 2007; Gibson 2009; Strömberg 2011; Bennett et al. 2013; Visser et al. 2013; Kellogg 2015; Linder et al. 2018).

**Figure 1.**
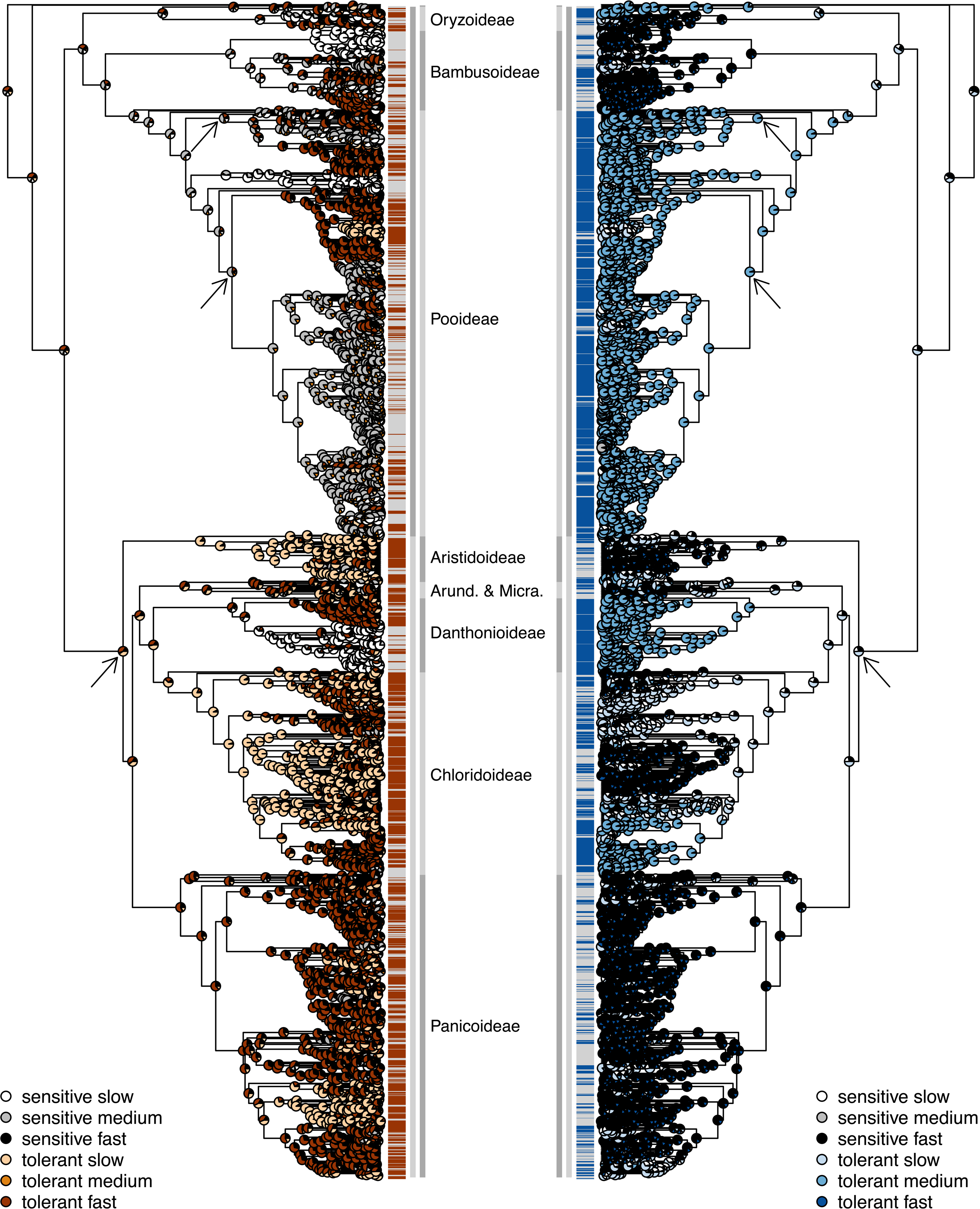
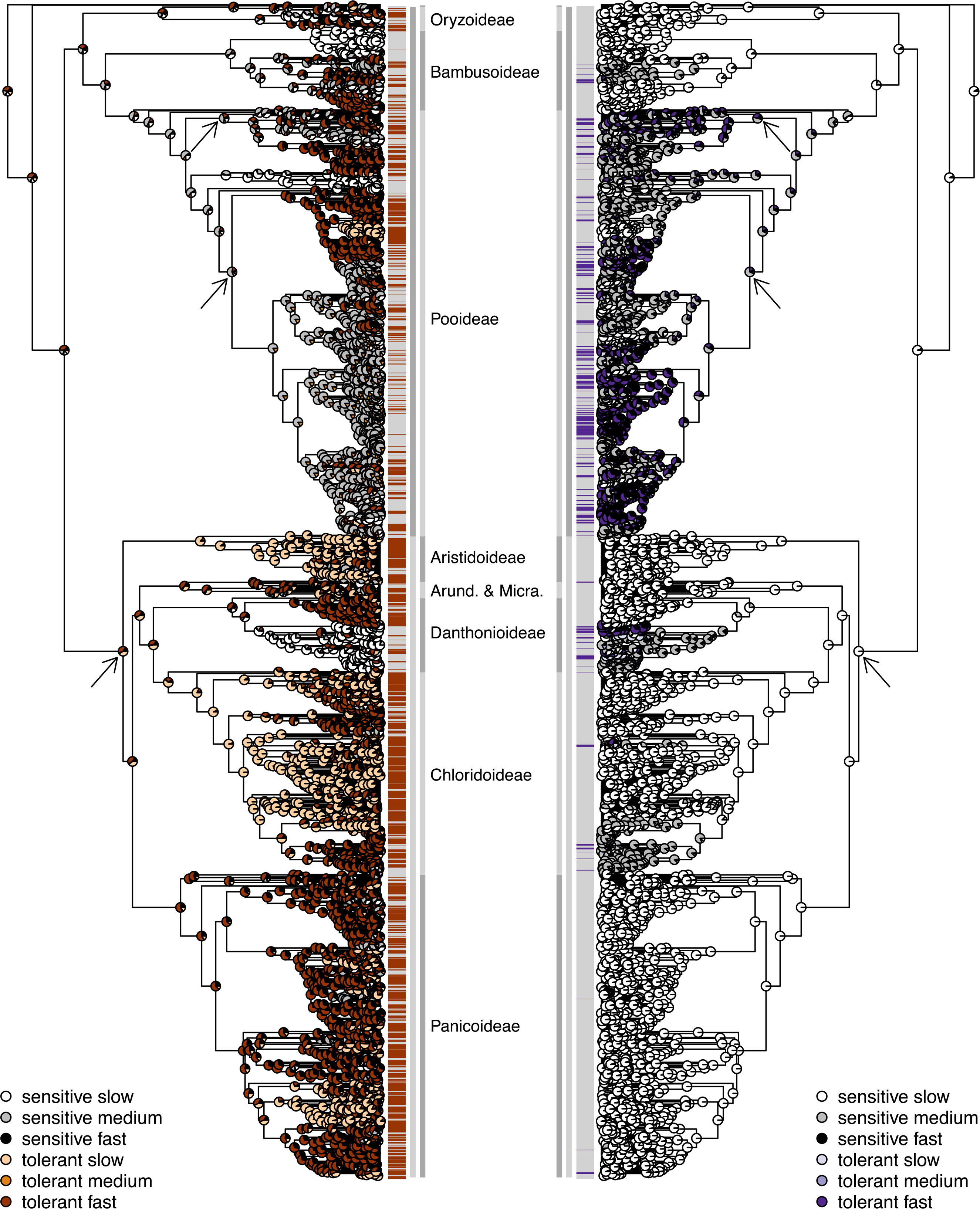
Ancestral state reconstructions for drought, frost and severe winter tolerance, displayed on facing trees for (a) drought-frost and (b) drought-winter. Ancestral states were inferred from the best-fitting hidden rates model (Tables 1 and 2). Pie charts indicate the marginal probabilities of the most likely state and rate at each node (the darker the shade, the higher the transition rate). Drought and frost tolerance in different rate categories are interpreted as representing different “types” of tolerance, corresponding to warm/cool climate drought tolerance and periodic/episodic frost tolerance, respectively (Supporting Text). Tolerance/sensitivity of the sampled species is labelled at the tips (red = drought tolerant, blue = frost tolerant, purple = winter tolerant, grey = sensitive). Outer grey vertical line: Subfamilies (“Arund. & Micra.” = Arundinoideae and Micrairoideae). Inner grey vertical line: the BOP clade (dark grey; Bambusoideae, Oryzoideae, Pooideae) and the PACMAD clade (light grey; Panicoideae, Aristidoideae, Chloridoideae, Micrairoideae, Arundinoideae, Danthonioideae). The two successive sisters to the rest sampled here (*Puelia olyriformis* and *Pharus latifolius*) are not part of any labelled subfamily or clade. Arrows indicate ancestral nodes where transitions from closed to open habitats may have occurred (Kellogg 2001; Bouchenak-Khelladi et al. 2010; Zhang et al. 2022; Elliott et al. 2023).

**Table 1:** Likelihood and AICc scores for ancestral state reconstructions with different numbers of hidden rate categories (CorHMM) for each of the three datasets^a^.

| Number of rate categories | Drought |  |  | Frost |  |  | Winter |  |  |
| --- | --- | --- | --- | --- | --- | --- | --- | --- | --- |
| 20% threshold | Log-likelihood | AICc | ΔAICc | Log-likelihood | AICc | ΔAICc | Log-likelihood | AICc | ΔAICc |
| One-rate | -1703.06 | 3410.12 | 340.06 | -1450.15 | 2904.31 | 415.81 | -891.76 | 1787.53 | 258.88 |
| Two-rate | -1542.86 | 3097.75 | 27.69 | -1262.47 | 2536.98 | 48.48 | -778.66 | 1569.36 | 40.71 |
| Three-rate | -1523.09 | 3070.30 | 0.24 | -1232.19 | 2488.50 | 0 | -752.27 | 1528.65 | 0 |
| Four-rate | -1514.88 | 3070.06 | 0 | -1229.63 | 2499.56 | 11.06 | -745.91 | 1532.13 | 3.48 |
| Five-rate | -1511.31 | 3083.30 | 13.24 | -1227.03 | 2514.74 | 26.24 | -743.61 | 1547.90 | 19.25 |
| **5% threshold** |  |  |  |  |  |  |  |  |  |
| One-rate | -1479.34 | 2962.68 | 347.24 | -1311.25 | 2626.50 | 355.91 | -1116.55 | 2237.10 | 326.34 |
| Two-rate | -1329.05 | 2670.13 | 54.69 | -1143.39 | 2298.80 | 28.21 | -977.41 | 1966.85 | 56.09 |
| Three-rate | -1295.66 | 2615.44 | 0 | -1123.21 | 2270.59 | 0 | -943.41 | 1910.93 | 0.17 |
| Four-rate | -1289.51 | 2619.32 | 3.88 | -1120.57 | 2281.45 | 10.86 | -935.23 | 1910.76 | 0 |
| Five-rate | -1288.19 | 2637.06 | 21.62 | -1120.32 | 2301.31 | 30.72 | -931.14 | 1922.96 | 12.20 |
| **50% threshold** |  |  |  |  |  |  |  |  |  |
| One-rate | -1629.85 | 3263.70 | 317.52 | -1388.07 | 2780.17 | 388.47 | -581.44 | 1166.89 | 165.37 |
| Two-rate | -1467.73 | 2947.50 | 1.32 | -1208.63 | 2433.32 | 41.62 | -499.67 | 1015.40 | 13.88 |
| Three-rate | -1461.04 | 2946.18 | 0 | -1185.64 | 2399.43 | 7.73 | -488.70 | 1001.52 | 0 |
| Four-rate | -1456.42 | 2953.14 | 6.96 | -1175.70 | 2391.70 | 0 | -482.10 | 1004.51 | 2.99 |
| Five-rate | -1454.33 | 2969.34 | 23.16 | -1171.75 | 2404.18 | 12.48 | -481.73 | 1024.13 | 22.61 |

<sup>a</sup>Shading denotes the best-fitting models ( ΔAICc<2; Burnham & Anderson, 2002).

**Table 2:** Rate matrix for the best-fitting hidden rates models (three-rate model) for the 20% threshold dataset^a^.

**a) Drought tolerance**
|  | Rate | Fast (R1) |  | Medium (R2) |  | Slow (R3) |  |
| --- | --- | --- | --- | --- | --- | --- | --- |
| Rate | Tolerance | 0 | 1 | 0 | 1 | 0 | 1 |
| Fast (R1) | 0 | - | 100.00 | 0.02 | - | 0.03 | - |
|  | 1 | 40.76 | - | - | 0.02 | - | 0.03 |
| Medium (R2) | 0 | 0.03 | - | - | 7.81 | 0.02 | - |
|  | 1 | - | 0.03 | 40.76 | - | - | 0.02 |
| Slow (R3) | 0 | 0.04 | - | 0.00 | - | - | 0.00 |
|  | 1 | - | 0.04 | - | 0.00 | 0.00 | - |

**b) Frost tolerance**
|  | Rate | Fast (R1) |  | Slow (R2) |  | Medium (R3) |  |
| --- | --- | --- | --- | --- | --- | --- | --- |
| Rate | Tolerance | 0 | 1 | 0 | 1 | 0 | 1 |
| Fast (R1) | 0 | - | 2.77 | 0.02 | - | 0.00 | - |
|  | 1 | 13.21 | - | - | 0.02 | - | 0.00 |
| Slow (R2) | 0 | 0.04 | - | - | 1.78 | 0.02 | - |
|  | 1 | - | 0.04 | 0.77 | - | - | 0.02 |
| Medium (R3) | 0 | 0.00 | - | 0.01 | - | - | 8.78 |
|  | 1 | - | 0.00 | - | 0.01 | 0.42 | - |

**c) Winter tolerance**
|  | Rate | Fast (R1) |  | Medium (R2) |  | Slow (R3) |  |
| --- | --- | --- | --- | --- | --- | --- | --- |
| Rate | Tolerance | 0 | 1 | 0 | 1 | 0 | 1 |
| Fast (R1) | 0 | - | 100.00 | 0.09 | - | 0.02 | - |
|  | 1 | 50.66 | - | - | 0.09 | - | 0.02 |
| Medium (R2) | 0 | 0.03 | - | - | 1.35 | 0.02 | - |
|  | 1 | - | 0.03 | 14.28 | - | - | 0.02 |
| Slow (R3) | 0 | 0.00 | - | 0.01 | - | - | 0.00 |
|  | 1 | - | 0.00 | - | 0.01 | 0.02 | - |

<sup>a</sup>Rates have been rounded off to two decimal places

Grasses are thought to have originated in the Cretaceous (ca. 93–125 Ma ago; Huang et al. 2016; Gallaher et al. 2019; Schubert et al. 2020), most likely in deep shade, as understorey plants of moist, warm forests (Bouchenak-Khelladi et al., 2010; Gallaher et al., 2019; Kellogg, 2001; Strömberg, 2011). Such habitats are today characteristic of a few small lineages that are successively sister to the rest of the family (Anomochloideae, Pharoideae and Puelloideae), the bamboos (Bambusoideae) and *Brachyelytrum*, which is sister to the rest of the Pooideae. However, the vast majority of grasses today occupy open environments (Elliott et al., 2023).

Transitions from closed to open environments are thought to have happened multiple times, certainly independently in the BOP and PACMAD clades, but exactly how many times and in which lineages is still unclear. Estimates range from one to as many as four independent transitions in the BOP clade, mainly in Pooideae, and one or possibly two in the PACMAD clade (Bouchenak-Khelladi et al., 2010; Elliott et al., 2023; Kellogg, 2001; Zhang et al., 2022). Both fossil and phylogenetic evidence suggests grasses began transitioning to open habitats around 67–58 Ma (Gallaher et al., 2019; Schubert, Marcussen, et al., 2019; Strömberg, 2011), at different times on different continents (Edwards et al., 2010; Strömberg, 2011). These transitions most lilkely occurred in response to increased aridity combined with altered disturbance regimes (seasonality, fire and herbivory), with less support for a role for global cooling (Edwards et al., 2010; Kellogg, 2015; Strömberg, 2011).

Similarly, grasses have transitioned from tropical to temperate climates multiple times independently, most notably leading to the radiation of two distantly related lineages into cool temperate zones, the Pooideae and Danthonioideae. However, frost tolerance has never been reconstructed as deeply as the nodes at which transitions to open habitats are implied (Edwards & Smith, 2010; Elliott et al., 2023; Humphreys & Linder, 2013; Linder et al., 2018; Sandve & Fjellheim, 2010; Schubert, Grønvold, et al., 2019). This suggests that grasses adapted to aridity before cold and frost and sets up predictions for which clades may have been predisposed to adapting to low temperatures under the hypothesis tested here. The level of cold and drought experienced by present-day grasses varies tremendously, e.g., with certain bamboos and PACMAD grasses also occurring in cold climates and cold temperate grasses occurring in both mesic (e.g. *Glyceria*) and dry (e.g. *Triticum*) habitats (Edwards & Smith, 2010; Humphreys & Linder, 2013; Schubert et al., 2020; Schubert, Marcussen, et al., 2019; Visser et al., 2013; Watcharamongkol et al., 2018; Zhang et al., 2022). Thus, the hypothesis that drought tolerance was a precursor trait to frost tolerance requires detailed analysis.

We use occurrence in areas that experience frost, severe winter and drought, according to the Köppen Geiger climate classification (Beck et al., 2018; Köppen, 2011), as a proxy for tolerance of each of these stresses. We refer to these stress tolerances in their broadest sense, e.g. ‘frost tolerance’ includes frost escape (e.g. spring annuals), frost avoidance or resistance (e.g. supercooling) and true frost tolerance (e.g. control of ice crystal formation). We then use phylogenetic comparative methods to test for evidence consistent with drought tolerance as a precursor for adapting to a temperate climate, specifically, to frost and severe winters. We first reconstruct ancestral states for drought, frost and winter tolerance separately to visually assess whether 1) frost and/or winter tolerant clades are nested within drought tolerant clades, or appear to arise more frequently in clades that were ancestrally drought tolerant. Next, because evolutionary transitions happen along branches and not at nodes, we model evolutionary transitions between sensitive and tolerant states simultaneously for two traits to test whether 2) drought tolerance evolves first and transitions to frost and winter tolerance are more frequent in drought tolerant than drought sensitive lineages. Finally, we 3) verify the models of correlated evolution with alternative null models and simulated data, and 4) test the effect of different species distribution thresholds for scoring species as drought, frost or winter tolerant.

## Methods

### Geographical and phylogenetic data

Geographical data were obtained from the Global Biodiversity Information Facility (GBIF, www.gbif.org). All GBIF entries with the family name “Poaceae” were downloaded on 25-02-2020 (https://doi.org/10.15468/dl.26b2o8), resulting in over 28 million observations. To remove erroneous and inaccurate observations, the following filtering steps were then followed. Observations that had coordinates with fewer than two decimals in either the latitude or longitude were removed. The CoordinateCleaner package (v. 2.0-15; Zizka et al., 2019) in R (v 3.6.3; R Core Team, 2020) was used to filter out observations with invalid coordinates, with equal or zero latitude and longitude coordinates, observations located in oceans, biodiversity institutions, GBIF headquarters, capital cities, country or province centroids and observations where the coordinates did not match the country code. Several major crop and horticultural species were removed manually, as these are likely to occur far outside their native ranges (and potentially climate niches). These were: wheat (*Triticum aestivum* L.), barley (*Hordeum vulgare* L.), sugarcane (*Saccharum officinarum* L.), corn (*Zea mays* L.), oats (*Avena sativa* L.), rice (*Oryza sativa* L.), sorghum (*Sorghum bicolor* (L.) Moench), rye (*Secale cereal* L.), ryegrasses (*Lolium multiflorum* Lam. and *Lolium perenne* L.), silvergrasses (*Miscanthus sinensis* Andersson and *Miscanthus giganteas* J.M.Greef & Deuter ex Hodk. & Renvoize), pampas grass (*Cortaderia selloana* (Schult. & Schult.f.) Asch. & Graebn.), switchgrass (*Panicum virgatum* L.) and several millets (*Setaria italica* (L.) P.Beauv.*, Eleusine coracana* (L.) Gaertn., *Cenchrus americanus* (L.) Morrone and *Panicum dichotomiflorum* Michx.).

Next, the geographical observations were matched to a phylogenetic tree comprising of 3595 species (Spriggs et al. 2014), using both the currently accepted species name (“species”) and the name used in the original GBIF entry (“verbatimScientificName”). This was done to minimise the chance of omitting species due to synonymy differences (e.g. if two species were previously considered separate but have now been synonymised, and are represented by two tips in the tree). We chose this method as currently synonymous tips were not always each other’s closest relatives, and we did not want to make (arbitrary) decisions on pruning one of these tips. However, this approach also opens the possibility of a single GBIF observation being matched to two different tips in the tree; however, this was true for only a tiny fraction of all data: in the final dataset 19,924 observations (0.33%) were matched to two species (tips). Overall, 2531 species with GBIF data could be matched to the tree using currently accepted names and 2546 using the name in the original GIBF entry. In the end, the filtering and species matching steps resulted in a dataset comprising of phylogenetic information and at least ten GBIF observations for 2731 species.

As part of our filtering process, we deemed geographical information based on fewer than ten GBIF observations per species unreliable and those species were therefore removed from the dataset (428 species). However, low taxon sampling may be at least as big of a limitation as few geographical occurrence records, and recent studies have included fewer or less precise geographical occurrence data (Pironon et al., 2024; Smith et al., 2023). To include as many species as possible, we therefore retained species with 5-9 GBIF observations if they are also known only from a single botanical country or first-level province, as defined under the World Geographical Scheme for Recording Plant Distributions by the Taxonomic Databases Working Group (TDWG; Brummitt et al. 2001).

Information on the distribution of grass species among TDWG regions was obtained from GrassBase (Clayton et al. 2006 onwards). This resulted in the inclusion of an additional 69 species, considered endemic (Supplementary Text, Supplementary Table 1), bringing the final total to 2800 species (Supplementary Figure 1). The number of geographical occurrence records per species in this dataset ranged from 5 to just over 1 million (for *Holcus lanatus* L. and *Dactylis glomerata* L.).

Finally, to prevent a bias when linking the geographical data to climate data due to high sampling in certain areas, and to remove duplicate GBIF entries, the geographical dataset was reduced to one observation per tile (in a 30 arc-second grid) per species using the raster package (v. 3.1-5; Hijmans, 2020). Following this, the number of unique observations per species ranged from 1 to 175,810, averaging at 2152 unique observations per species. The final dataset consisted of 2800 species with both geographical and phylogenetic data and just over 6 million GBIF observations.

### Defining areas experiencing drought, frost and severe winter using Köppen-Geiger climate zones

It is notoriously difficult to define drought in an ecologically meaningful way, and to delimit areas that experience drought (Slette et al., 2019). Because of this, we used Köppen-Geiger climate zones to define geographical areas experiencing drought, frost and severe winter (Supplementary Table 2). Köppen-Geiger zones divide the world into five main climate zones based on their vegetation composition: Tropical (*A*), Arid (*B*), Temperate (*C*), Continental (*D*) and Polar (*E)*, and each zone is then further divided into subzones based on temperature and precipitation criteria (Köppen 2011; Beck et al. 2018). Each zone was scored as experiencing drought, frost and/or winter as follows. Zones *B* (arid and semi-arid climates), *Aw* and *As* (tropical savannahs) were scored as experiencing drought. Additionally, zone *Csa* was scored as experiencing drought due to its combination of high temperatures and low precipitation during the summer months (e.g. Los Angeles, Perth and Valencia).

For consistency, we also used Köppen-Geiger zones to define areas experiencing frost and severe winters (Supplementary Table 2). Köppen-Geiger zones classified as experiencing frost were defined as having one or more months averaging below 0 °C; the *D* and *E* zones fulfilled this criterion. Although zones *Bxk*, *Cxb*, and *Cxc* do not have any months averaging below 0 °C, they do experience occasional to regular frost during the winter months (e.g. Reno [Nevada, USA] and the Patagonian Desert for *Bxk*; Cape Town, Mexico City and Plymouth [UK] for *Cxb*; and Balmaceda [Chile], Bodø [Norway] and the Faroe Islands for *Cxc*). Therefore, they were also scored as experiencing frost. Zones with severe winter were defined as those experiencing frost and a cool, short growing season with a maximum of three months averaging above 10 °C. This included zones *Cxc*, *Dxc*, *Dxd* and *E* (Supplementary Table 2).

### Species distribution thresholds and sensitivity analyses

Following the scoring of which geographical areas experience drought, frost and severe winter, the GBIF geographical occurrence records were linked to this scoring to determine which species are exposed to drought, frost or severe winter in their native ranges using the R packages kgc (v. 1.0.0.2; Bryant et al., 2017) and raster (v. 3.1-5; Hijmans, 2020). We acknowledge that inferring physiological tolerances from occurrence patterns can both underestimate and overestimate true tolerances, e.g. if species distributions are strongly structured according to microclimatic variation (Greiser et al., 2020). However, it is still a useful approach at macroevolutionary scales (Humphreys & Linder, 2013).

We considered species drought, frost and/or winter tolerant if at least 20% of their geographical observations fell in Köppen-Geiger zones scored accordingly. We chose this threshold to remove species occurring, but not established, in areas with drought, frost or severe winter, with an emphasis on representing each species’ optimal climate niche rather than its extremes. These species will hereafter be referred to as “drought tolerant”, “frost tolerant” and “(severe) winter tolerant”, respectively. We also tested whether our results are contingent on the use of this threshold by creating two alternative datasets in which species were considered “tolerant” if at least 5% or 50% of their occurrence records fell in an area experiencing drought, frost and/or winter. Analyses were run on data compiled using all three thresholds (20%, the “main” dataset; 5% and 50%, the “sensitivity” datasets), making nine datasets in total (3 traits *x* 3 thresholds).

### Ancestral state reconstructions of drought, frost, and winter tolerance

To visualise the pattern of evolution for each of drought, frost and winter tolerance separately, we performed ancestral state reconstructions using the “hidden rates” Markov models implemented in the R package corHMM (Beaulieu et al., 2013, 2020). This method takes rate heterogeneity during the course of evolution and among clades into account, and has been shown to perform similarly to other rate-variable methods (King & Lee, 2015). In the simplest corHMM model, the one-rate category model, a binary trait evolves with the same rate across the entire tree. This is equivalent to a standard model for a binary trait with one rate of gain and one rate of loss (all rates different; Beaulieu et al., 2013). The constraints on the model are then relaxed, stepwise, to generate increasingly complex models by adding additional rate categories. The two-rate category model thus allows each state to evolve in a fast and a slow rate category, with each rate category having its own rate of both gain and loss, as well as a rate of moving to the other rate category. In the three-rate category model, there are fast, medium, and slow rate categories, and so on, until the most complex model currently implemented in corHMM, the five-rate category model (Beaulieu et al., 2013). The models do not allow for both the trait and transition rate category to change simultaneously (e.g. from “sensitive-slow” to “tolerant-fast”), but multiple shifts of either type can happen along a single branch. The rate of changing from one rate category to another is independent of the trait (e.g. the rate of moving from the “fast” to the “slow” rate category is the same irrespective of whether the state is “sensitive” or “tolerant”). We fitted models with different numbers of transition rate categories, with the states “tolerant” and “sensitive”, across all nine datasets using maximum likelihood (ML) criteria and 100 random restarts. Model fit was assessed using Akaike’s sample-size corrected information criterion (Burnham & Anderson, 2002; AICc). The estimated parameters of the best-fitting model were used to calculate the marginal probabilities of the most likely tolerant and sensitive state and rate at each internal node. Most interpretations of model inferences were based on nodes with a marginal probability ≥0.75 for either tolerance or sensitivity (Beaulieu et al., 2013).

### Testing for correlated evolution between drought and frost/winter tolerance

To test for correlated evolution between drought tolerance and each of frost and winter tolerance, we tested whether the evolution of one trait (rate of gains and losses of frost and winter tolerance) is dependent on the state of the second trait (drought tolerance) by comparing the likelihood fit of two models (Pagel, 1994). The first model, the independent model, has four transition rate parameters, meaning that the rate of change in one trait is independent of the state of the second trait (Supplementary Figure 2a). The second model, the correlated model, has eight possible rate parameters, meaning that the rate of change in one trait may differ depending on the state of the second trait (Supplementary Figure 2b). If drought tolerance was a precursor for frost and/or winter tolerance, we would expect support for the correlated model, such that drought tolerance evolves first from the ancestral state (sensitive to drought, frost and severe winter) and transitions to frost and/or winter tolerance are more frequent in drought tolerant than drought sensitive lineages. We fitted the independent and correlated models using the one-rate category corHMM models as above for both trait combinations (drought-frost and drought-winter) and all three thresholds (six analyses in total). Drought-winter tolerance based on the 50% dataset was tested against a reduced model (Supplementary Text, Supplementary Figure 2c). Model fit was assessed using AICc.

As with several macroevolutionary methods, models of correlated evolution have been shown to have high type I error rates, i.e., rejection of a null hypothesis that is true (Boyko & Beaulieu, 2023; Humphreys et al., 2016; Maddison & FitzJohn, 2015; Rabosky & Goldberg, 2015). There can be several underlying causes of this, so we addressed this in two ways:

#### I. Testing for correlated evolution using a null model with rate heterogeneity

False rejection of the null model may be because it is too trivial to serve as a meaningful null. This has been suggested for the independent model used here, specifically, because it fails to take rate heterogeneity into account (Boyko & Beaulieu, 2023). Thus, we compared the model of correlated evolution to an alternative null model: the independent model with two rate categories (the two-rate model). This model is formulated as two paired independent models, with four transition rates in each of the two rate categories (Supplementary Figure 2d). Additionally, there are two transition rates for transitioning between rate categories. Thus, the two-rate independent model has ten rate parameters. We fitted the two-rate model across all six trait and threshold combinations using corHMM as before (Supplementary Text, Supplementary Figures 2d,e). Model fit was assessed using AICc.

#### II. Testing for correlated evolution using simulated data

False rejection of the null model may be due to it not adequately explaining the evolution of the traits over the tree – but may be unrelated to any hypothesised correlation between the two traits. Therefore, we compared support for the correlated and independent models using data simulated to represent a binary trait that has evolved independently of either frost or winter tolerance. We expect that the null (independent) model should not be rejected in favour of the alternative (correlated) model for simulated data. If it is, then the difference in fit between the correlated and independent models needs to be significantly smaller for simulated than observed (empirical) data for any observed correlation to remain statistically significant.

We simulated 1000 continuous traits on the observed phylogenetic tree under Brownian motion, using ‘bmPlot’ in the R package phytools (Revell, 2012). Trait values were drawn from a normal distribution centred around zero. Each trait was then converted to a binary trait with the same proportions of sensitive and tolerant species as for the observed drought tolerance data (20% threshold; Supplementary Table S3), by scoring the 1549 species with the highest values as tolerant, and the 1251 remaining species as sensitive.

Independent and correlated models with one transition rate category were then fitted to each simulated trait plus frost and winter tolerance in turn (20% threshold), using the Discrete independent and Discrete dependent functions in BayesTraits (available from www.evolution.rdg.ac.uk; Pagel et al. 2004). First, exploratory models were fitted using 1000 ML iterations. Next, the models were fitted using a Bayesian version of the method (Pagel et al., 2004). The models were run for 1 million Markov chain Monte Carlo (MCMC) generations, all with 1000 stones with 10,000 iterations each using default settings. Model support was determined using Bayes Factors (BFs).

## Results

### Distribution of sampled species and occurrence records among clades and Köppen-Geiger climate zones

Grasses have been recorded from every Köppen-Geiger zone. Of all filtered GBIF observations, 64% occurred in the *Cfb* zone (temperate oceanic climate), which covers large parts of Western Europe, New Zealand and South-East Australia. The zones with the second and third most records were *Cfa* (humid subtropical climate) and *Dfc* (subarctic climate), accounting for only 5.8% and 5.5% of the observations, respectively. Of the 2800 species in the dataset, 1549 species occur in areas experiencing drought, 1759 species occur in areas experiencing frost, 375 species occur in areas with severe winters, 828 species are exposed to both drought and frost and 57 species to both drought and severe winter (20% threshold dataset; Supplementary Table 3). The phylogeny includes around 25% of all Poaceae species, but species sampling is unevenly distributed among subfamilies (Supplementary Text, Supplementary Table 4), and GBIF records are unevenly distributed among species (Supplementary Text, Supplementary Figure 3).

### Ancestral state reconstructions

We report the results for the 20% threshold dataset. Results for the 5% and 50% threshold datasets are summarized below. Under the 20% threshold dataset, *Pharus latifolius* L., sister to all other species sampled here, is scored as sensitive to drought, frost and severe winter (Figure 1).

#### Drought tolerance

All major clades contain species scored as drought tolerant (Figure 1), however, three clades stand out as having fewer drought tolerant species: Pooideae (specifically Poeae), Bambusoideae (specifically Arundinarieae) and Danthonioideae (the clade sister to *Pentameris*). The two best-fitting models for drought tolerance were the three-rate and four-rate models (Λ1AICc ≥ 13.24), which were statistically indistinguishable from each other (Λ1AICc = 0.24; Table 1). Therefore, we accepted the simpler three-rate model. Under this model, the ancestors of the PACMAD clade and all its constituent subfamilies were most likely drought tolerant (Figure 1), with the highest marginal probabilities (>=0.75) for ancestral drought tolerance for PACMAD as a whole and all subfamilies except Panicoideae and Danthonioideae (Supplementary Figure S4). However, several clades within Panicoideae (e.g. subtribe Boivinellinae) and Danthonioideae were reconstructed as being ancestrally drought tolerant (Supplementary Figure S4). The reconstruction for the ancestors of the BOP clade and Poaceae as a whole was unccertain (Figure 1, Supplementary Figure 4). Within the BOP clade, no deeper nodes were reconstructed as being drought tolerant, but certain clades within Pooideae (e.g. *Aegilops* and within Stipeae), Bambusoideae and Oryzoideae were reconstructed as drought tolerant. The ancestors of Pooideae (except *Brachyelytrum* that is sister to the rest) and Bambusoideae were most likely drought sensitive, as was the ancestor of Oryzoideae but with less certainty (Figure 1, Supplementary Figure 4).

Based on the sum of the gains and losses of tolerance, we identified a fast (R1), medium (R2), and slow (R3) rate category (Table 2). Drought tolerance in the fast and slow rate categories are reconstructed throughout the tree (Figure 1): the fast category is reconstructed in all major clades, notably in much of Panicoideae, half of Danthonioideae (*Pentameris*), parts of Chloridoideae, Bambusoideae, Oryzoideae and parts of Pooideae (tribes Stipeae and Triticeae); the slow rate category is primarily reconstructed in the Aristidoideae, Chloridoideae, a small number of lineages in Panicoideae (e.g. subtribe Boivinellinae), and *Aegilops* (Triticae, Pooideae). Drought sensitivity is inferred mainly in the medium (R2) and slow (R3) rate categories, with the medium category being prevalent in the Pooideae (especially Poeae) and the slow category reconstructed in Bambusoideae (Arundinarieae) and Danthonioideae (sister clade to *Pentameris*).

#### Frost tolerance

Species belonging to all subfamilies were scored as being frost tolerant, with the Danthonioideae, Chloridoideae and Pooideae having the most frost tolerant species (Figure 1a). The three-rate category model was the best fit (ΔAICc ≥ 11.06; Table 1). Under this model, reconstructed ancestral states deeper in the tree are equivocal, but more certain for some of the subfamilies. Pooideae, Danthonioideae and part of Chloridoideae (*Muhlenbergiinae sensu* Peterson et al. 2010) were most likely ancestrally frost tolerant and Bambusoideae and Panicoideae (excluding two small clades successively sister to the rest) most likely ancestrally frost sensitive (marginal probability >= 0.75; Figure 1a, Supplementary Figure 4a).

Based on the sum of gains and losses, we distinguish fast (R1), medium (R3) and slow (R2) rate categories (Table 2). Frost tolerance in the medium rate category is inferred in the Pooideae, Oryzoideae (*Ehrharta*), Danthonioideae and Chloridoideae (*Muhlenbergiinae sensu* Peterson et al. 2010; Figure 1a). Frost tolerance in the slow category is less frequently occurring and distributed more sporadically throughout the tree, notably being inferred in the Chloridoideae (*Zoysieae*, *Eragrostidinae*) and Panicoideae. Frost tolerance in the slow category is more easily lost than frost tolerance in the medium category (relative rate of loss 0.77 compared to 0.42, Table 2). For frost sensitive clades, the fast rate category (R1) is most common but the slow category is often reconstructed equivocally at the same nodes as frost tolerance in the slow category.

*Severe winter tolerance* – Species scored as tolerant of severe winter are clustered mainly in Pooideae and Danthonioideae, but a few winter tolerant species occur also in other subfamilies (Figure 1b). The three-rate category model had the best fit for winter tolerance (ΔAICc ≥ 3; Table 1). Poaceae as a whole, the BOP and PACMAD clades and each of the subfamilies were all inferred as being ancestrally winter sensitive (marginal probability >=0.75; Figure 1b, Supplementary Figure 4b). Very few internal nodes were reconstructed as winter tolerant, the exceptions being scattered clades within Danthonioideae (e.g *Chionochloa*) and Pooideae (in Poeae, Triticeae and Stipeae; marginal probability >=0.75; Figure 1b, Supplementary Figure 4b).

Based on the sum of gains and losses, we define the rate categories fast (R1), medium (R2) and slow (R3; Table 2). Winter tolerance is mostly found in the fast rate category (Figure 1b). This winter tolerance is often associated with, and evolves from, winter sensitivity in the medium and fast rate categories but never in the slow rate category. Lineages in the medium rate sensitive state are found in Pooideae and parts of Danthonioideae (sister to *Pentameris*) but also, notably, in *Muhlenbergiinae sensu* Peterson et al. 2010 (Chloridoideae) and the neotropical woody bamboos (Bambusoideae). Most other clades are not tolerant of severe winters and ancestral states are inferred to be sensitive in the slow category.

### Models of correlated evolution between drought and frost/winter tolerance

The correlated model was a better fit than the independent model for both drought-frost and drought-winter (ΔAICc > 59; Table 3). However, the two-rate independent model was much better than either of the single-rate models, for both drought-frost and drought-winter (ΔAICc > 234; Table 3). Both ML and Bayesian analyses of simulated data and frost tolerance revealed that the difference in fit between the independent and correlated models for the simulated data overlapped with the difference in fit between the two models for the observed data (Figures 2a,b). For simulated data and winter tolerance, the difference in fit between the correlated and independent models was greater in the observed compared to simulated data (Figures 2b,c; the observed BF was outside the 95% confidence of intervals of BFs for the simulated data). However, the nature of the correlation inferred from the best-fitting model was higher rates of gains of winter tolerance in drought sensitive than tolerant lineages.

**Figure 2.**
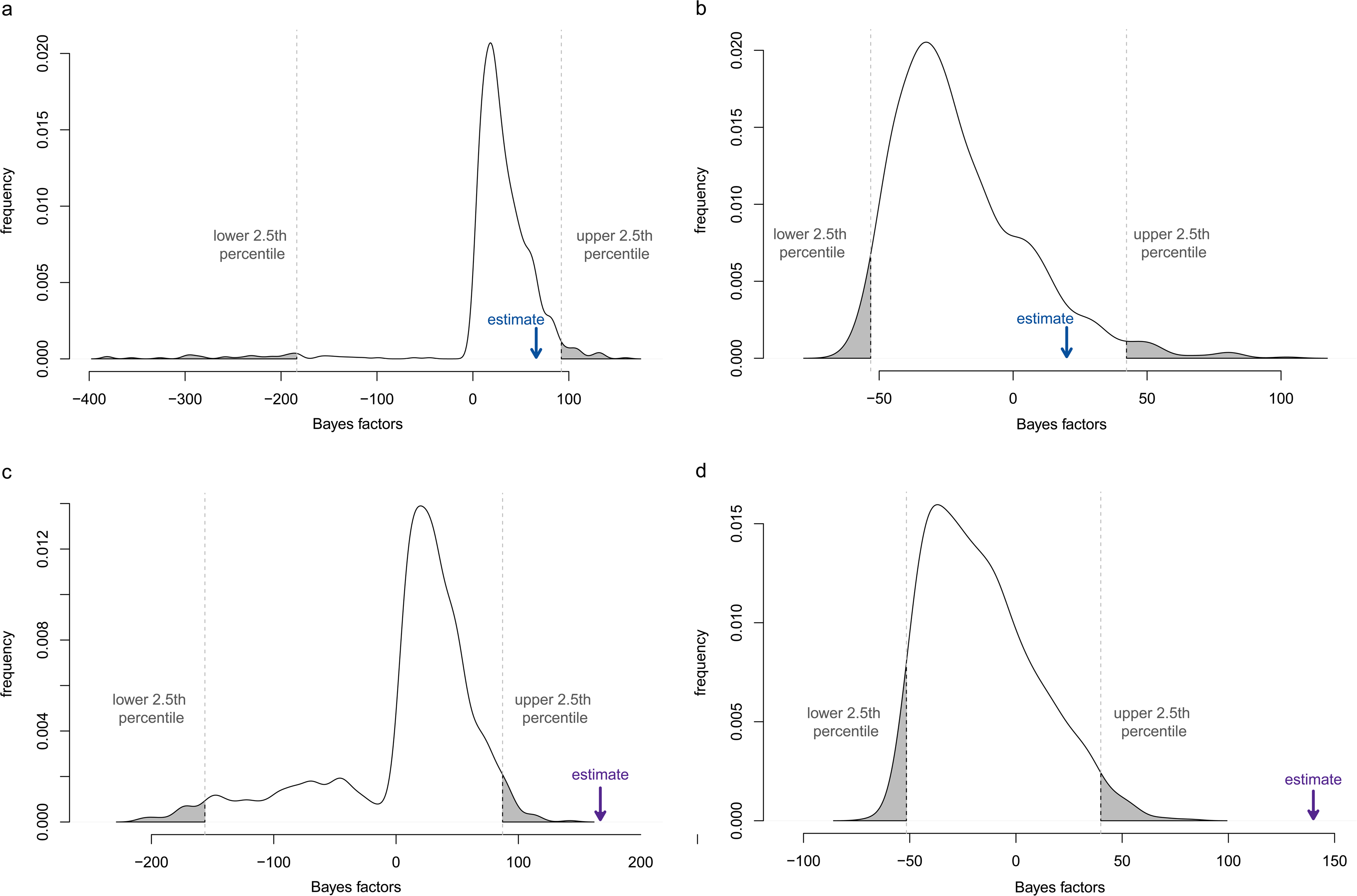
Difference in fit between independent and correlated models of evolution. (a,b) Models fitted to 1000 simulated traits and observed frost tolerance data using maximum likelihood (a) and Bayesian (b) methods. (c,d) Models fitted to 1000 simulated traits and observed winter tolerance data using maximum likelihood (c) and Bayesian (d) methods. The difference-in-fit 95% confidence intervals (CI) for simulated data are indicated by vertical, dashed lines, with the area outside the CI shaded. Arrows indicate the difference in fit for the observed drought-frost (a,b) and drought-winter (c,d) data.

**Table 3:**
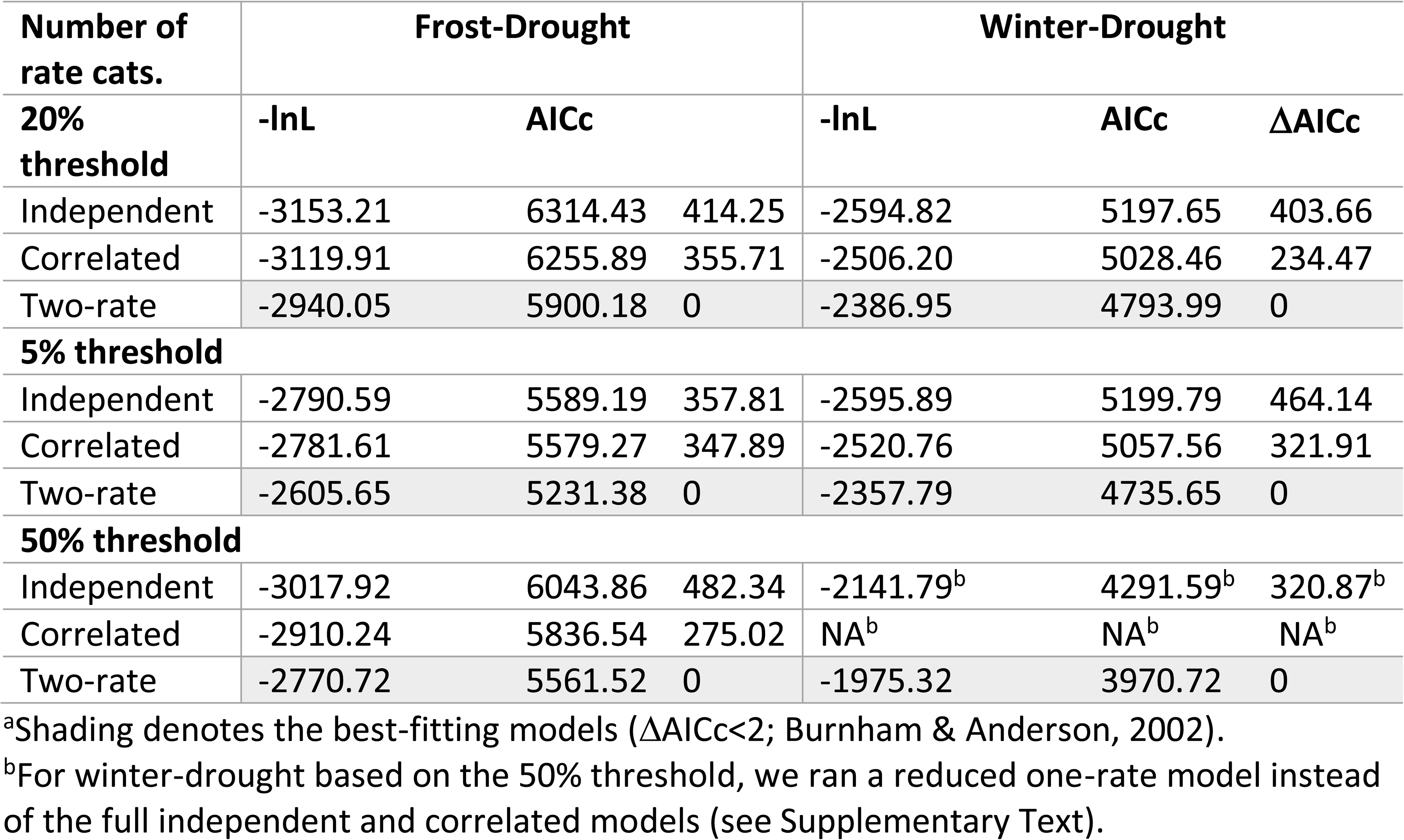
Likelihood and AICc scores for the one-rate independent, one-rate correlated, and two-rate independent models for each of the three datasets^a^.

### Sensitivity analyses

The main results were the same across all datasets, with six exceptions (Supplementary Text; Tables 1, 3, Supplementary Table 3). Four differences were for the 50% threshold datasets and two were for the 5% threshold datasets. The differences concerned scoring of drought tolerance for *Pharus latifolius*, scoring of severe winter tolerance in Bambusoideae and model fit. None of these differences affected the overall reconstructed evolutionary scenario underlying the conclusions.

## Discussion

We tested for a signature of evolutionary precursors in adaptation to a temperate climate. The importance of precursors in the formation of biomes and assembly of regional floras is known in the biogeographical literature as ‘biome conservatism’ (Crisp et al., 2009) or the ‘temperate element’ of high alpine floras (Gehrke and Linder 2009; Gehrke 2018; but see Nürk et al. 2018). Biome transitions require complex structural, physiological and phenological adjustments, and it is reasonable to assume that not all such changes happened simultaneously. Instead, it is more plausible that present-day adaptations were acquired gradually, with the gradual accumulation of multiple new traits providing opportunities for persisting in one changing environment, as well as colonising other emerging environments.

Despite some research (e.g. Sakai and Larcher 1987; Judd et al. 1994; Zanne et al. 2014; Spriggs et al. 2015; Edwards et al. 2017), little is known, however, about the order in which adaptations to freezing and seasonal conditions originated in present-day temperate clades. Little is also known about the extent to which individual lineages still hold signatures of any ancient precursor traits that may have facilitated their biome transitioning journey. To address this, we used a comparative approach to test the idea that tropical-to-temperate transitions were facilitated by precursor trait(s). Specifically, we tested for evidence that drought tolerance acted as a precursor to frost and winter tolerance in grasses. That is, that drought tolerance evolved first and frost and winter tolerance originated more frequently in ancestrally drought tolerant lineages, suggesting that ancestral drought tolerance facilitated transitions into cold temperate environments (Preston & Sandve, 2013; Sakai & Larcher, 1987).

### Ancestral state reconstructions reveal both drought-first and frost-first scenarios

Drought tolerance is inferred to have evolved early in the PACMAD clade, with the ancestor most likely being drought tolerant (Figure 1; Supplementary Figure 4). This is consistent with drought tolerance evolving simultaneously with the transition from closed to open environments in the ancestor of the PACMAD clade (Elliott et al., 2023; Kellogg, 2001; Strömberg, 2011). In contrast, drought tolerance was not reconstructed deep in the BOP clade. This is consistent with multiple independent transitions to open habitats in the BOP clade, although our reconstruction of drought tolerance suggests it evolved long after transitions to open environments occurred (Elliott et al., 2023; Kellogg, 2001; Zhang et al., 2022). The relatively late evolution of drought tolerance in Pooideae is consistent with the “tippy” distribution of xerophytes and high evolutionary lability of drought responses found previously for this clade (Pal Stolsmo et al., 2024; Zhang et al., 2022).

Overall, our reconstruction suggested a later origin of frost tolerance than drought tolerance (Figure 1a; Supplementary Figure 4a). Frost tolerance was not inferred as ancestral in either the BOP or PACMAD clades, but the ancestors of Pooideae, Danthonioideae and part of Chloridoideae (*Muhlenbergiinae*; Peterson et al. 2018) were reconstructed as frost tolerant. The overall deeper origin of drought than frost tolerance supports the view that grasses were exposed to drought before frost during their evolutionary history (Linder et al., 2018; Strömberg, 2011). Deep origins of frost tolerance in Pooideae and Danthonioideae is also consistent with previous studies (Humphreys & Linder, 2013; Schubert, Marcussen, et al., 2019). However, upon scrutiny, it is clear that frost tolerant lineages in the BOP and PACMAD clades have undergone different evolutionary trajectories; drought tolerance most likely preceded frost tolerance in Danthonioideae and Chloridoideae but not in Pooideae. In other words, frost tolerance evolved in ancestrally drought tolerant lineages in Danthonioideae and Chloridoideae, but in ancestrally drought sensitive lineages in Pooideae.

Tolerance of severe winters was reconstructed ancestrally for certain clades within Danthonioideae (especially *Chionochloa*) and Pooideae (within tribes Poeae, Triticeae and Stipeae; Figure 1b). In addition, scattered occurrences of winter tolerance were found in Bambusoideae, Chloridoideae and Panicoideae (Figure 1b). In all these clades except Panicoideae, winter tolerance is preceded by sensitivity in the medium rate category, suggesting that this state may represent a precursor trait (Figure 1b; but see Supplementary Text for the reconstruction in Bambusoideae). Tolerance of severe winters encompasses both physiological and phenological adaptations, and a plausible prerequisite is adaptation to periodic (seasonal) frost. Our results corroborate this, at least for Pooideae and parts of Danthonioideae and Chloridoideae (Supplementary Text, Supplementary Figure 5). Since we found patterns consistent with drought tolerance as a precursor to frost tolerance in the latter two clades, drought tolerance could indirectly be a precursor to severe winter tolerance in those clades as well.

### Models of correlated evolution unsupported – with important caveats

The above discussion is based on comparison of separate analyses for each of drought, frost and winter tolerance. The explicitly simultaneous modelling of drought-frost and drought-winter tolerance showed that the initial rejection of the model of independent evolution in favour of the correlated model is due to a type I error. This is because: 1) the correlated model was not a better fit than the alternative null model that allowed for rate heterogeneity (Table 3); and 2) the difference in fit between the correlated and independent models for the observed frost-drought data overlapped with the difference in fit for data simulated under independent evolution (Figure 2). For drought-winter, the difference in fit was significantly greater for the observed than simulated data but the best-fitting correlated model suggested that winter tolerance evolved in drought sensitive lineages, not drought tolerant ones (not shown). Taken at face value, these results imply that evolution of drought and frost tolerances were not correlated and that drought tolerance is unlikely to have acted as a precursor to either frost or winter tolerance.

However, there are two caveats to this interpretation. The first is technical. The two-rate model fitted here is formulated as two paired independent models with a total of ten transition rates (Supplementary Figure 2d). Increasing model complexity to fully address the question of correlated evolution while adequately accounting for rate heterogeneity (Boyko & Beaulieu, 2023; King & Lee, 2015) would likely improve model fit, but such models would soon become prohibitively complex and computationally demanding for a dataset of this size. This means we are unlikely to have found the best models for evolution of drought-frost or drought-winter, we have only demonstrated that the initial rejection of the independent model was likely due to statistical error.

The second caveat is biological. Our ancestral state reconstructions reveal different evolutionary trajectories in different clades (cf. the BOP and PACMAD clades, Figure 1, Supplementary Figure 4). Modelling the Poaceae as a whole may have obscured these different trajectories. A straightforward solution would be to analyse the BOP and PACMAD clades separately; however, our results point to a single deep origin of drought tolerance in the PACMAD clade, followed by repeated origins of frost tolerance. This scenario thus reduces to a single evolutionary event that lacks the comparison of ancestrally drought sensitive clades (Maddison & FitzJohn, 2015; Uyeda et al., 2018). This leaves the option of addressing the question at even broader phylogenetic scales (e.g. the Poales), but this approach is marred by the challenge of accurately modelling the many diverse histories that are inevitable at such broad scales. Thus, in lieu of more complex modelling in carefully selected clades (Donoghue & Edwards, 2014), our conclusion based on separate ancestral state reconstructions for each trait separately is a drought-first scenario in the PACMAD clade but a frost-first scenario in the BOP clade.

## Conclusions

Two different evolutionary trajectories are evident from our results. Drought tolerance most likely evolved first in the PACMAD clade, followed by repeated entries into freezing climates, followed in Danthonioideae by adaptation to severe winters. In the BOP clade, frost tolerance most likely evolved first, followed by adaptation to severe winters in some clades and dry environments in other clades. The drought-first scenario for the PACMAD clade is consistent with our expectations and evidence that grasses were exposed to dry before freezing conditions (e.g. Strömberg 2011; Linder et al. 2018). In light of this, the frost-first scenario for the BOP clade is surprising, but consistent with recent evidence from ecophysiological experiments and gene expression patterns in Pooideae (Das et al., 2023; Pal Stolsmo et al., 2024), and suggestions that Pooideae originated in emerging cool climate pockets present as early as the Late Cretaceous (Das et al., 2023; Hagen et al., 2019; Schubert et al., 2020). Therefore, it is likely that frost tolerance evolved before drought tolerance in the BOP clade. Alternatively, high evolutionarily lability of drought tolerance (Table 2; Pal Stolsmo et al. 2024) limits the accuracy with which we can reconstruct how cold and drought traits evolved relative to each other in this clade (Bromham, 2015). Thus, although it is likely that modern-day stress responses originated from shared ancient pathways (Das et al., 2023; Folk et al., 2020; Preston & Sandve, 2013), it is possible that the signature of those pathways is no longer be present in some clades.

The different reconstructed evolutionary trajectories reveal a mirrored pattern, with clades occupying either predominantly dry or very cold environments. This mirrored pattern can be seen throughout the grasses (*cf.* the two main clades of Danthonioideae (*Pentameris* vs. its sister clade), several clades in Pooideae (e.g. Stipeae, *Aegilops* and *Triticum*), and the PACMAD versus BOP clades; Figure 1, Supplementary Figure 4), and is consistent with modern-day grasses being either drought or cold specialists (Visser et al., 2013). This suggests a trade-off, such that species tend not to be equally well adapted to both drought and cold, as has been found for woody temperate plants (Puglielli et al., 2022).

Finally, what is the consequence of relying on geographical occurrences instead of inherent physiological tolerances? While the relative tolerances of species are likely related to their distributions (Humphreys & Linder, 2013), absence from a region experiencing drought, frost or severe winter, as defined here, does not necessarily equate to an inability to withstand those stresses. As such, our method may be underestimating the prevalence of each adaptation across the family (Humphreys & Linder, 2013; Zanne et al., 2014). Further, if species distributions follow microclimatic variation, our reliance on coarse-grid occurrence records could both underestimate and overestimate the prevalence of drought and frost tolerant species. Severe winter tolerance, as defined here, should be less affected by microclimatic specialisations as seasonal cold and frost coupled with short growing seasons occur across large, high-latitude areas. Overall, however, our reconstructions captured the patterns seen in previous ancestral state reconstrucitons for grasses (e.g. Edwards & Smith, 2010; Elliott et al., 2023; Humphreys & Linder, 2013; Schubert et al., 2020; Schubert, Marcussen, et al., 2019; Watcharamongkol et al., 2018; Zhang et al., 2022), are robust to alternative treatments of the data (Supplementary Text) and corroborate findings for Pooideae, based on measured ecophysiological responses to frost and drought exposure (Pal Stolsmo et al., 2024). Future research addressing these concerns is needed but they are unlikely to be biasing our results here.

## Supporting information

Schatetaal_Drought Tolerance as a Percursor_2024_Supplementary Material

## Author contributions

L.S., M.S., S.F. and A.M.H. designed the study. L.S. compiled the data and performed the analyses, and all authors contributed to the interpretation of the results. L.S. and A.M.H. wrote the manuscript with input from M.S. and S.F.

## Funding

This research has been funded by the Carl Trygger Foundation (CTS 18:156) and the Bolin Centre for Climate Research, Stockholm University.

## Conflict of interest statement

The authors declare no conflict of interest.

## Acknowledgements

We are grateful to B. Gehrke for fruitful discussions, and we thank L. Gren for assistance with putting together Figure 2.

## Notes

### Competing Interest Statement

The authors have declared no competing interest.

