## Supplementary material for "Drought tolerance as an evolutionary precursor to frost and winter tolerance in grasses": Schatetaal_Drought Tolerance as a Percursor_2024_Supplementary Material

### Supplementary Text

#### Sensitivity analyses

*Methods*

To test the dependency of our empirical results on the use of the 20% threshold for distinguishing tolerant from sensitive species, all analyses were run on two more datasets: one in which species were scored as tolerant if at least 5% of their geographical occurrences were in Köppen-Geiger climate zones experiencing drought, frost and/or winter and one in which species were scored as tolerant if at least 50% of their occurrence records fell in these areas (Supporting Table; Beck et al., 2018; Köppen, 2011). The 50% threshold dataset contains no species that are both drought and winter tolerant (Supplementary Table 3). Therefore, to test for correlated evolution between drought and winter tolerance in this dataset, we created reduced models, by removing the option of transitioning to and from the state of being both drought and winter tolerant (cf. Boyko & Beaulieu, 2023). The reduced model in the one-rate category had four rate parameters (Supplementary Figure 2c), and therefore we could not distinguish between the dependent and independent models. The reduced two-rate model had 10 rate parameters (Supplementary Figure 2e). We compared the fit of the reduced one-rate and two-rate models for the 50% threshold dataset for drought-winter tolerance.

*Results*

The main results were the same across all datasets, with six exceptions (Tables 1,3, Supplementary Table 3). Two differences were for the 5% threshold datasets. Firstly, *Pharus* *latifolius*, which is sister to all other species sampled here and thus assumed to be drought sensitive, was scored as drought tolerant with the 5% threshold but drought sensitive with the 20% threshold. Several other species were scored differently for all three traits (Supplementary Table 3) but this had a negligible effect on downstream analyses for the 5% threshold datasets. The only difference for these datasets was for the hidden rates analyses for winter tolerance, where the three- and four-rate category models had similar fits ($\Delta$AICc = 0.17; Table 1), whereas the three-rate category model was better for the 20% dataset. The simpler three-rate category model revealed the same pattern of reconstructed ancestral states as for the 20% threshold (Figures 1, S4 for the 20% threshold; not shown for the 5% threshold). This difference was therefore minor and overall, there were no significant differences between the 5% and 20% threshold datasets.

Several species were scored differently under the 50% threshold for all three traits (Supplementary Table 3) but there were only four differences pertaining model fit and inferences. Firstly, for drought tolerance, the two- and three-rate category models had similar fits ($\Delta$AICc = 1.32; Table 1), whereas the three-rate model was supported for the 20% dataset. The simpler two-rate category model revealed a similar pattern of reconstructed ancestral states as the 20% threshold dataset (not shown). This difference is therefore negligible. Secondly, for frost tolerance, the four-rate model was the best fit (Table 1), whereas the three-rate model was supported for the 20% dataset. However, this model also yielded the same ancestral state reconstruction overall (not shown) and is therefore minor. Thirdly, for winter tolerance, the three-rate category model was best-fitting, as it was for the 20% threshold dataset (Table 1). However, the reconstructed scenario differed for the neotropical woody bamboos (Bambusoideae), which were inferred as winter sensitive in the medium rate category for the 20% threshold and winter sensitive in the slow rate category for the 5% threshold (not shown). This is caused by the different scoring of species between the two datasets, as there are no winter tolerant species based on the 50% threshold (cf. Figure 1b). All ancestral states in the bamboos are inferred as winter sensitive (>= 75% of ML iterations; Supplementary Figure 4b) but any interpretation of the rate category of that sensitivity may be unreliable. Finally, we were not able to fit the one-rate correlated and independent models for drought-winter tolerance based on the 50% threshold dataset because there are no species that are both drought and winter tolerant. Comparison of the reduced one-rate and two-rate models (Supplementary Figure 2c,e) revealed that the two-rate model was the best fit (Table 3), which is in agreement with the results for the 20% threshold dataset. Thus, overall, the effects of the differing species distribution thresholds compared here are minor, except for determining the rate category of ancestral winter sensitivity in Bambusoideae.

*Discussion*

The overall proportions of frost tolerant species differ slightly from previous studies but patterns across clades remain consistent (Humphreys & Linder, 2013; Watcharamongkol et al., 2018; Schubert et al., 2019, 2020). Differences in the estimated proportions of frost tolerant species may be explained by the different methods employed for defining frost tolerant species. We used Köppen-Geiger climate zones to define freezing areas and then scored species based on a given proportion of their occurrence records that fell in those areas. However, choosing a threshold value (i.e. proportion of occurrences) for distinguishing tolerance from sensitivity is a largely arbitrary decision – is presence in freezing/winter conditions enough to make a species tolerant, or does a certain amount of a species' range need to overlap with freezing areas (Edwards et al., 2015; Zanne et al., 2014)? This issue is not trivial and can have significant consequences for the biological conclusions we draw, especially if poorly known species are affected (Bätscher & de Vos, 2024; Cousins-Westerberg et al., 2023; Edwards et al., 2015). In this study, results were largely robust to the choice of threshold value. Even so, we consider the 5% and 50% threshold datasets somewhat extreme and therefore would expect them to be prone to error (e.g. leading to an overestimation/underestimation of drought/frost/winter tolerance in Poaceae). More generally, however, we cannot see that there is a universal 'best' threshold for scoring species as tolerant or sensitive to a particular climatic condition. Our findings emphasise the importance of testing multiple thresholds, while also considering the number of observations per species, the trait and the question of interest.

**Distribution of sampled species and occurrence records among clades and Köppen-Geiger climate zones**

The data used for our analyses are unevenly distributed among clades and regions, displaying patterns that largely mirror known sampling biases in biodiversity data. For example, the phylogenetic tree (based on Spriggs et al. 2014) contains approximately 25% of all described Poaceae species (WCSP; Brummitt et al., 2001; GrassBase; Clayton et al., 2006); however, this percentage differs among subfamilies, being highest for Danthonioideae (63%) and lowest for Bambusoideae (13%; Supplementary Table 4). In addition, Bambusoideae and *Pentameris* (Danthonioideae) had the lowest numbers of GBIF observations per species (ranging 1 to ca. 50

records after the filtering and reduction steps), while several species in Pooideae were represented by >1000 observations (Supplementary Figure 3). These patterns confirm the phylogenetic biases found previously for New England grasses (Daru et al. 2018) and may have affected our results for Bambusoideae. This clade is inferred to possess a putative precursor for severe winter tolerance (sensitivity in the medium rate category) but, in contrast to the other clades with this state, is inferred to have episodic rather than periodic frost tolerance (see below under '*Two types of drought and frost tolerance inferred*'). However, this result is not robust to the sensitivity analyses (see above) and undersampling may have led to an underrepresentation of frost tolerant species in this clade.

The geographical collection bias apparent in our data, with the majority of the geographical occurrence records being from temperate oceanic climates (Köppen-Geiger zone *Cfb*), corresponds to that determined by Vorontsova et al. (2020), with grasses being overrepresented in temperate and seasonally dry tropical zones. These patterns also confirm a more general bias in biodiversity data towards more occurrence records in countries with high GDPs (Meyer et al., 2016) and Human Development Indexes (Vorontsova et al., 2020). Zone *Cfb* is scored as experiencing frost and the large quantity of data from this zone might have led to an overrepresentation of frost tolerant grasses, if, for example, a widespread species is highly sampled in zone *Cfb* but undersampled in non-freezing zones. Thus, frost/winter tolerant species might be overrepresented overall, or in certain clades, and underestimated in others (bamboos). By and large, however, our study captures known patterns in grasses (Figure 1), that are robust to different treatments of the data, as well as revealing new ones.

**Including species with few GBIF observations**

Preliminary analyses showed that including the 69 species with fewer than 10 GBIF observations in the analyses did not affect our findings. Previous studies have shown that species with few herbarium records are generally rare (Enquist et al., 2019), and rare species tend to have small geographic ranges and narrow niches (Rabinowitz, 1981). Thus, including species with little GBIF data, but which are known to be rare and/or narrow-ranged based on alternative data sources (e.g. TDWG for plants; Brummitt et al. 2001), is a promising approach for maximising species sampling in broadscale analyses (Pironon et al., 2024; Smith et al., 2023).

**Two types of drought and frost tolerance inferred**

We inferred drought tolerance in two different rate categories, suggesting the existence of two different "types" of drought tolerance in grasses. Drought tolerance in the fast rate category is reconstructed throughout the grasses, while the slow rate category is prevalent in clades occupying warm temperate and arid to semi-arid climates (KG zones BSh, BSk, BWh and Cfa), e.g. PACMAD clades with $C_4$ photosynthesis (Edwards & Smith, 2010; Watcharamongkol et al., 2018). A single instance of this type of drought tolerance was found in the BOP clade, in *Aegilops*, a genus of $C_3$ annuals primarily from warm, semi-arid climates (KG zones Csa and BSk; Kellogg, 2015). In contrast, the other type of drought tolerance (fast category) is found mainly in species from subtropical highlands and continental climates (KG zones D, Cfb and Cwb). These patterns suggest that the two types of drought tolerance correspond to occurrence in warm-arid versus cool-semi-arid areas, respectively. The extent to which this reflects different underlying physiologies and life history strategies warrants further research.

Our reconstruction also suggests two different rate categories of frost tolerance (Figure 1a), consistent with the existence of two "types" of frost tolerance. The first type (slow rate category) was reconstructed in clades that are mainly (sub)tropical and primarily experience occasional, short episodes of frost (e.g. Panicoideae and Chloridoideae, but also *Aegilops*, Pooideae). The second type (medium rate category) occurs in the cool temperate clades, Pooideae and Danthonioideae, which experience prolonged periods of frost, and also in *Muhlenbergia* from colder semi-arid climates (Chloridoideae; KG zone BSk) and *Ehrharta* from subtropical highlands (Oryzoideae; KG zone Cfb). Thus, these two types of frost tolerance could correspond to exposure to episodic (short-term, diurnal) and periodic (long-term, seasonal) frost, respectively. Episodic and periodic frost tolerance require different physiological adaptations and regulatory mechanisms, including cold acclimation ability and reduced photosynthetic activity and growth at low temperatures during periodic frost exposure (Schubert et al. 2020). Furthermore, even for those lineages inferred to be evolving in the same rate category (same type of frost tolerance), different physiological mechanisms are likely, since the limited evidence available suggests differing strategies of true frost tolerance in Pooideae compared to Danthonioideae (Humphreys & Linder, 2013; Schubert et al., 2020; Wharton et al., 2010). Further research is needed to determine the extent to which the two types of frost tolerance found here correspond to different underlying response mechanisms.

#### Supporting Table 1: Endemic species[a] included in dataset

| Species name | TDWG zone[b] | TDWG L3[b] | Species name | TDWG zone[b] | TDWG L3[b] |
|---|---|---|---|---|---|
| *Acidosasa notata* | China Southeast | CHS | *Muhlenbergia maxima* | Peru | PER |
| *Ampelocalamus microphyllus* | China South-Central | CHC | *Nassella cabrerae* | Argentina Northwest | AGW |
| *Apocopis courtallumensis* | India | IND | *Oligostachyum lubricum* | China Southeast | CHS |
| *Bambusa distegia* | China South-Central | CHC | *Panicum auricomum* | Brazil North | BZN |
| *Bonia amplexicaulis* | China Southeast | CHS | *Panicum cipoense* | Brazil Southeast | BZL |
| *Bromus striatus* | Peru | PER | *Panicum longipedicellatum* | Brazil South | BZS |
| *Capeochloa setacea* | Cape Provinces | CPP | *Panicum loreum* | Brazil Southeast | BZL |
| *Chimonocalamus delicatus* | China South-Central | CHC | *Panicum poliophyllum* | Brazil Southeast | BZL |
| *Chimonocalamus pallens* | China South-Central | CHC | *Panicum restingae* | Brazil Northeast | BZE |
| *Chionochloa spiralis* | New Zealand South | NZS | *Paspalum ramboi* | Brazil South | BZS |
| *Chusquea arachniformis* | Columbia | CLM | *Pentameris longiglumis* | Cape Provinces | CPP |
| *Chusquea nudiramea* | Brazil South | BZS | *Pentameris scandens* | Cape Provinces | CPP |
| *Cortaderia boliviensis* | Bolivia | BOL | *Pentameris swartbergensis* | Cape Provinces | CPP |
| *Danthonia annableae* | Bolivia | BOL | *Phalaris amethystina* | Chile Central | CLC |
| *Danthonia malacantha* | Chile Central | CLC | *Piptochaetium setosum* | Chile Central | CLC |
| *Digitaria catamarcensis* | Argentina Northwest | AGW | *Poa chathamica* | Chatham Islands | CTM |
| *Dinochloa scabrida* | Borneo | BOR | *Poa exigua* | New Zealand South | NZS |
| *Eriochloa setosa* | Cuba | CUB | *Poa lindebergii* | Krasnoyarsk | KRA |
| *Festuca cundinamarcae* | Columbia | CLM | *Poa xenica* | New Zealand South | NZS |
| *Festuca cuzcoensis* | Peru | PER | *Psathyrostachys caduca* | Afghanistan | AFG |
| *Festuca donax* | Madeira | MDR | *Pseudosasa orthotropa* | China Southeast | CHS |
| *Festuca flacca* | Ecuador | ECU | *Pseudosasa owatarii* | Japan | JAP |
| *Festuca fragilis* | Venezuela | VEN | *Racemobambos hepburnii* | Borneo | BOR |
| *Festuca luciarum* | New Zealand North | NZN | *Reederochloa eludens* | Mexico Northeast | MXE |
| *Festuca yalaensis* | Hawaii | HAW | *Sasa longiligulata* | China Southeast | CHS |
| *Germainia pilosa* | Thailand | THA | *Schizachyrium gaumeri* | Mexico Southeast | MXT |
| *Gigantochloa wrayi* | Malaya | MLY | *Setaria grandis* | Malawi | MLW |
| *Glaziophyton mirabile* | Brazil Southeast | BZL | *Stipa annua* | Peru | PER |
| *Hordeum erectifolium* | Argentina Northeast | AGE | *Stipa kingii* | California | CAL |
| *Hordeum guatemalense* | Guatemala | GUA | *Temburongia simplex* | Borneo | BOR |
| *Indosasa hispida* | China Southeast | CHS | *Trichoneura elegans* | Texas | TEX |
| *Ischaemum santapaui* | India | IND | *Trichoneura weberbaueri* | Peru | PER |
| *Lithachne humilis* | Honduras | HON | *Yushania baishanzuensis* | China Southeast | CHS |
| *Lycochloa avenacea* | Lebanon-Syria | LBS | *Zizaniopsis villanensis* | Argentina Northeast | AGE |

| | | |
|---|---|---|
| *Muhlenbergia involuta* | Texas | TEX |

[a]Species occurring in only a single TDWG level 3 region[2] with 5-9 observations following filtering.
[b]TDWG = Taxonomic Databases Working Group, a custom, standardized system of country and first-level province boundaries developed for plants (Brummitt et al., 2001). Level 3 regions refer to countries or first-level provinces.

**Supporting Table 2: Scoring of Köppen-Geiger climate zones for drought, frost and severe winter**

| | Zone[a] | Frost | Winter | Drought |
|---|---|---|---|---|
| **A (Tropical)** | Af | 0 | 0 | 0 |
| | Am | 0 | 0 | 0 |
| | As | 0 | 0 | 1 |
| | Aw | 0 | 0 | 1 |
| **B (Arid)** | Bsh | 0 | 0 | 1 |
| | Bsk | 1 | 0 | 1 |
| | Bwh | 0 | 0 | 1 |
| | Bwk | 1 | 0 | 1 |
| **C (Temperate)** | Cfa | 0 | 0 | 0 |
| | Cfb | 1 | 0 | 0 |
| | Cfc | 1 | 1 | 0 |
| | Csa | 0 | 0 | 1 |
| | Csb | 1 | 0 | 0 |
| | Csc | 1 | 1 | 0 |
| | Cwa | 0 | 0 | 0 |
| | Cwb | 1 | 0 | 0 |
| | Cwc | 1 | 1 | 0 |
| **D (Continental)** | Dfa | 1 | 0 | 0 |
| | Dfb | 1 | 0 | 0 |
| | Dfc | 1 | 1 | 0 |
| | Dfd | 1 | 1 | 0 |
| | Dsa | 1 | 0 | 0 |
| | Dsb | 1 | 0 | 0 |
| | Dsc | 1 | 1 | 0 |
| | Dsd | 1 | 1 | 0 |
| | Dwa | 1 | 0 | 0 |
| | Dwb | 1 | 0 | 0 |
| | Dwc | 1 | 1 | 0 |
| | Dwd | 1 | 1 | 0 |
| **E (Polar)** | Ef | 1 | 1 | 0 |
| | Et | 1 | 1 | 0 |

[a]Köppen-Geiger climate zones. These divide the world into five main climate zones based on their vegetation composition: Tropical (*A*), Arid (*B*), Temperate (*C*), Continental (*D*) and Polar (*E)*, and each zone is then further divided into subzones based on temperature and precipitation criteria (Köppen 2011; Beck et al. 2018).

**Supporting Table 3: Number of drought, frost and severe winter tolerant and sensitive species (and combinations thereof) for each of the three datasets**

| | **Drought Tolerant** | **Drought sensitive** | **Sum** |
|---|---|---|---|
| **20% threshold** | 1549 | 1251 | **2800** |
| Frost tolerant | 828 | 931 | 1759 |
| Frost sensitive | 721 | 320 | 1041 |
| Winter tolerant | 57 | 318 | 375 |
| Winter sensitive | 1492 | 933 | 2425 |
| **5% threshold** | 1974 | 826 | **2800** |
| Frost tolerant | 1457 | 686 | 2143 |
| Frost sensitive | 517 | 140 | 657 |
| Winter tolerant | 242 | 334 | 576 |
| Winter sensitive | 1732 | 492 | 2224 |
| **50% threshold** | 971 | 1829 | **2800** |
| Frost tolerant | 224 | 1100 | 1324 |
| Frost sensitive | 747 | 729 | 1476 |
| Winter tolerant | 0 | 191 | 191 |
| Winter sensitive | 971 | 1638 | 2609 |

| Supporting Table 4: Percentage of species in Poaceae and its subfamilies included in the tree | | | |
|---|---|---|---|
| **Subfamily** | **Species[a]** | **Species in tree** | **Sampling[a] %** |
| Aristidoideae | 365 | 110 | 30% |
| Arundinoideae and Micrairoideae | 243 | 39 | 17% |
| Bambusoideae | 1441 | 190 | 13% |
| Chloridoideae | 1721 | 484 | 28% |
| Danthonioideae | 281 | 177 | 63% |
| Oryzoideae | 112[b] | 59 | 53% |
| Panicoideae | 3316 | 722 | 22% |
| Pooideae | 3850 | 1017 | 26% |
| **Poaceae** | **11369[c]**<br>**11454[d]** | **2800** | **25%[c]**<br>**24%[d]** |

[a]Based on Kellogg (2015) unless specified otherwise
[b]Reported as Ehrhartoideae in Kellogg (2015)
[c]Based on GrassBase (Clayton et al., 2006; according to Vorontsova et al., 2020)
[d]Based on World Checklist of Vascular Plants (Brummitt et al., 2001; according to Vorontsova et al., 2020)

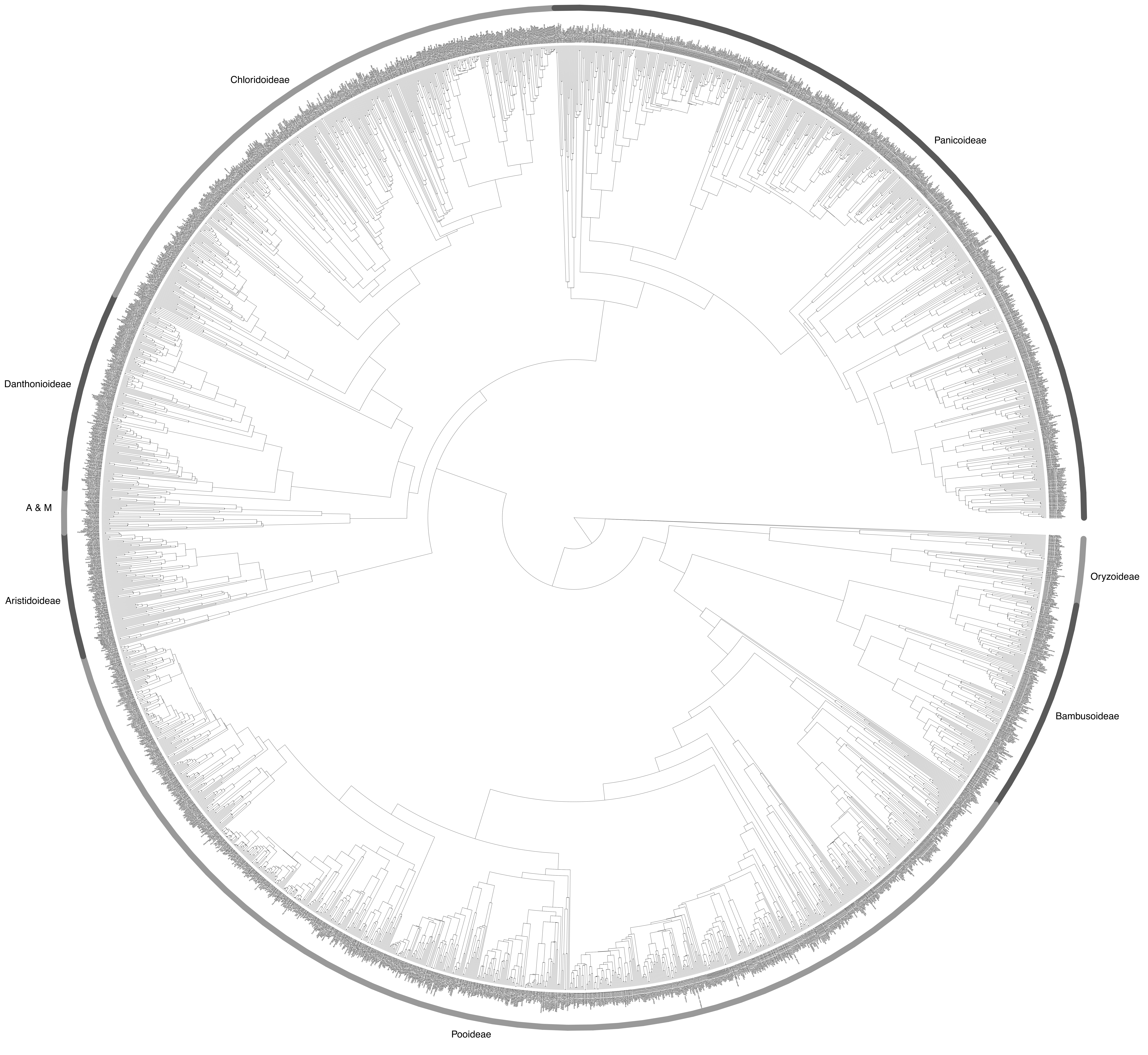
Chloridoideae
Panicoideae
Danthonioideae
A & M
Aristidoideae
Oryzoideae
Bambusoideae
Pooideae

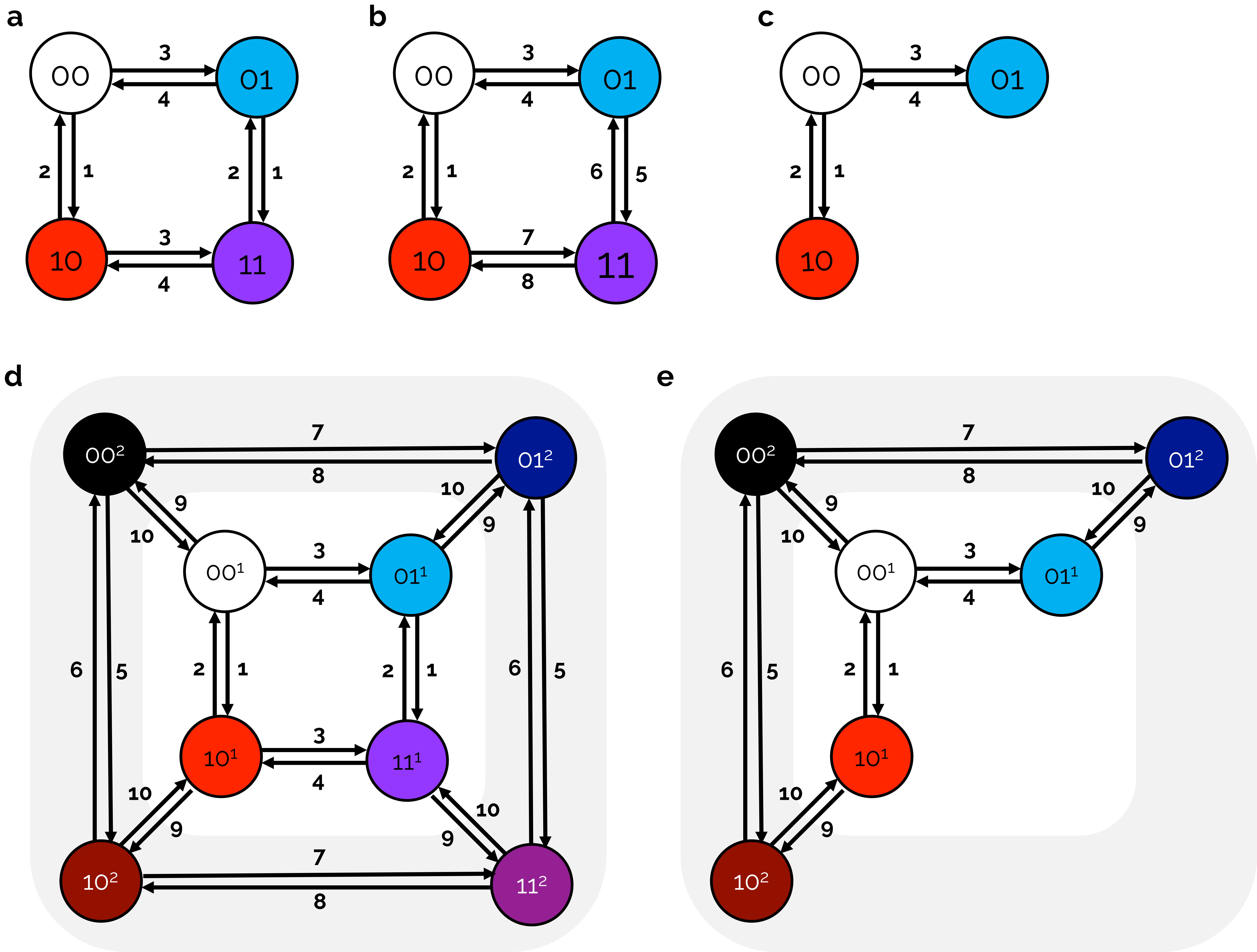
a
00
01
10
11
b
00
01
10
11
c
00
01
10
d
00²
01²
00¹
01¹
10¹
11¹
10²
11²
e
00²
01²
00¹
01¹
10¹
10²

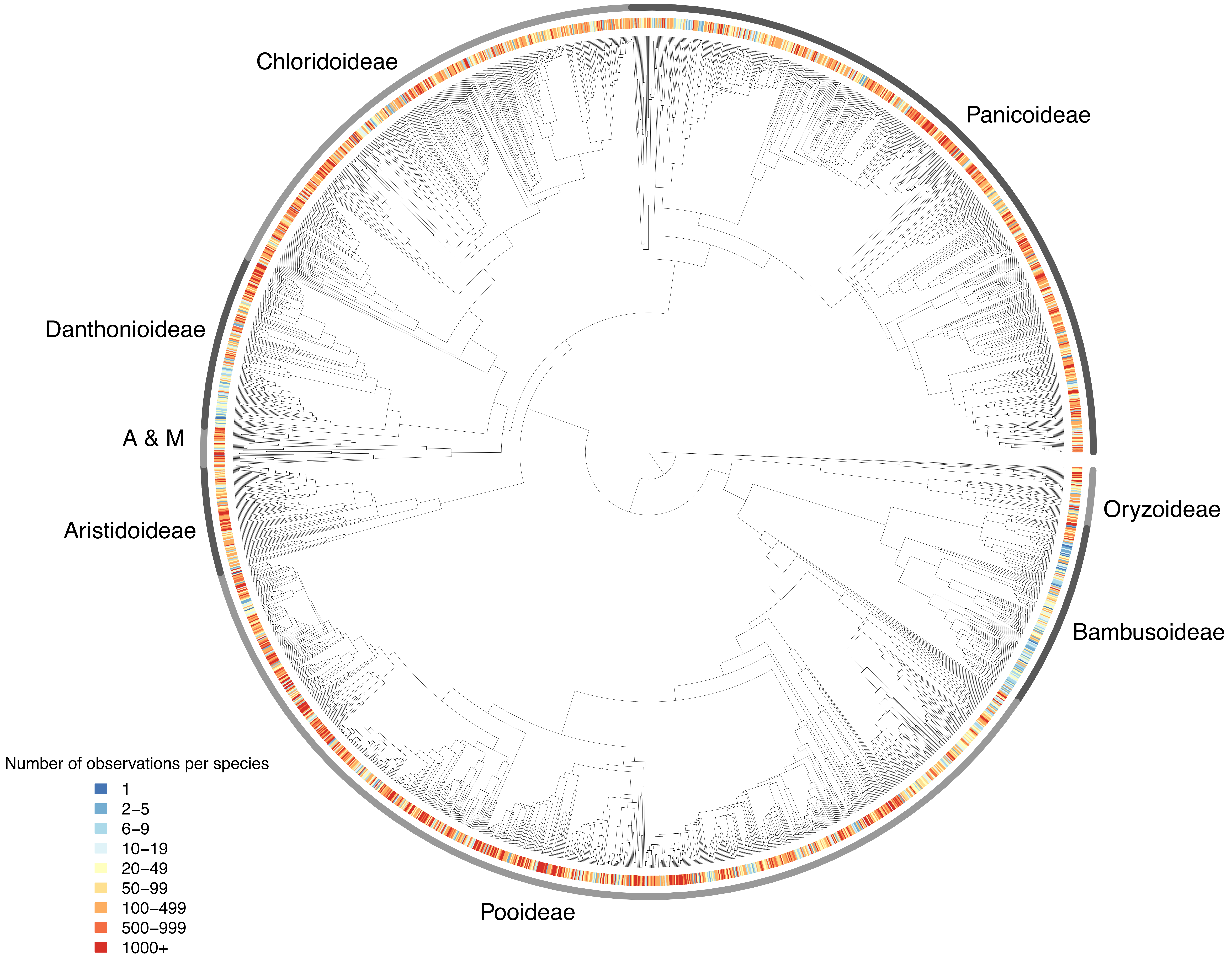
Chloridoideae
Panicoideae
Danthonioideae
A & M
Aristidoideae
Oryzoideae
Bambusoideae
Pooideae
Number of observations per species
1
2–5
6–9
10–19
20–49
50–99
100–499
500–999
1000+

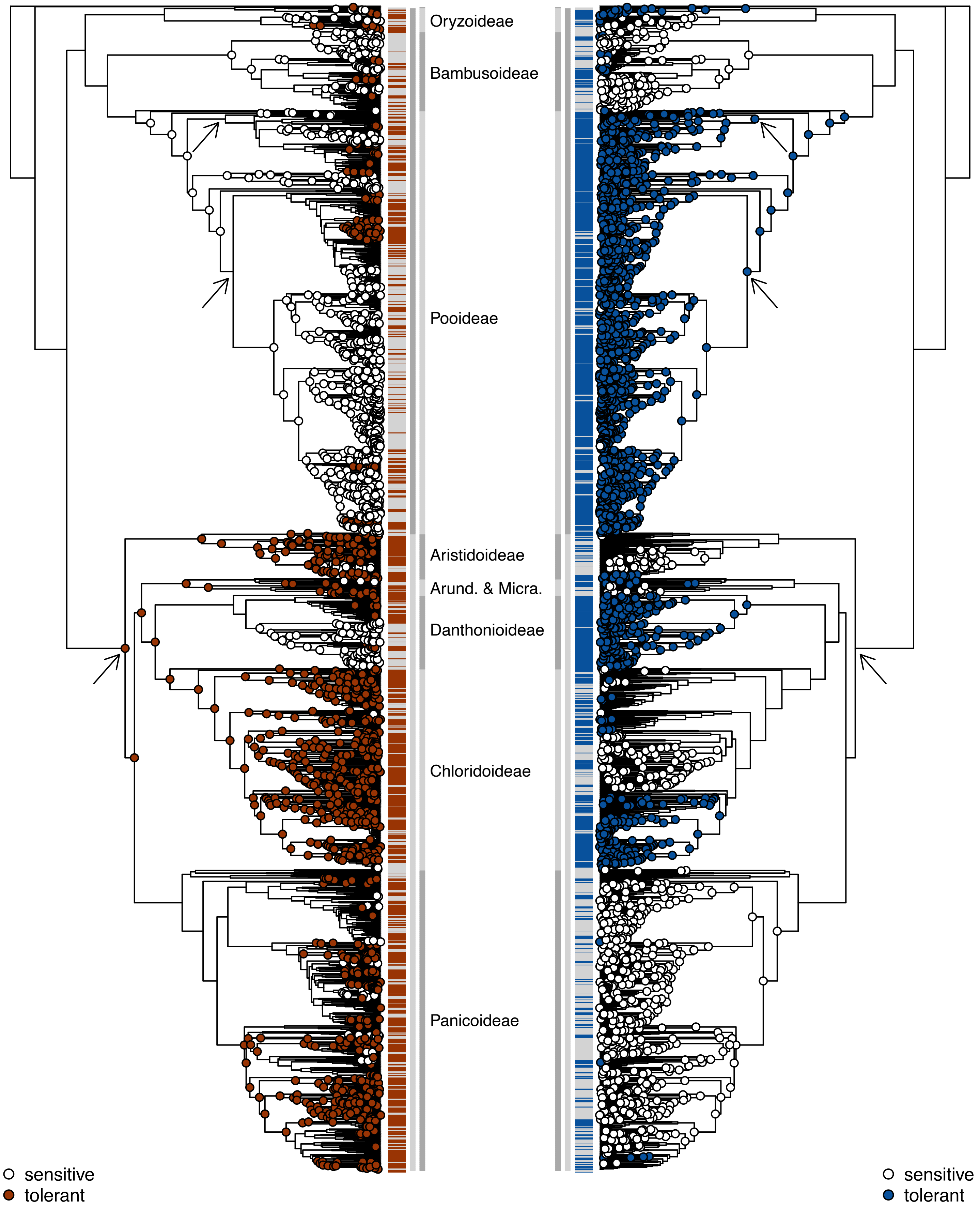
Oryzoideae
Bambusoideae
Pooideae
Aristidoideae
Arund. & Micra.
Danthonioideae
Chloridoideae
Panicoideae
sensitive
tolerant
sensitive
tolerant

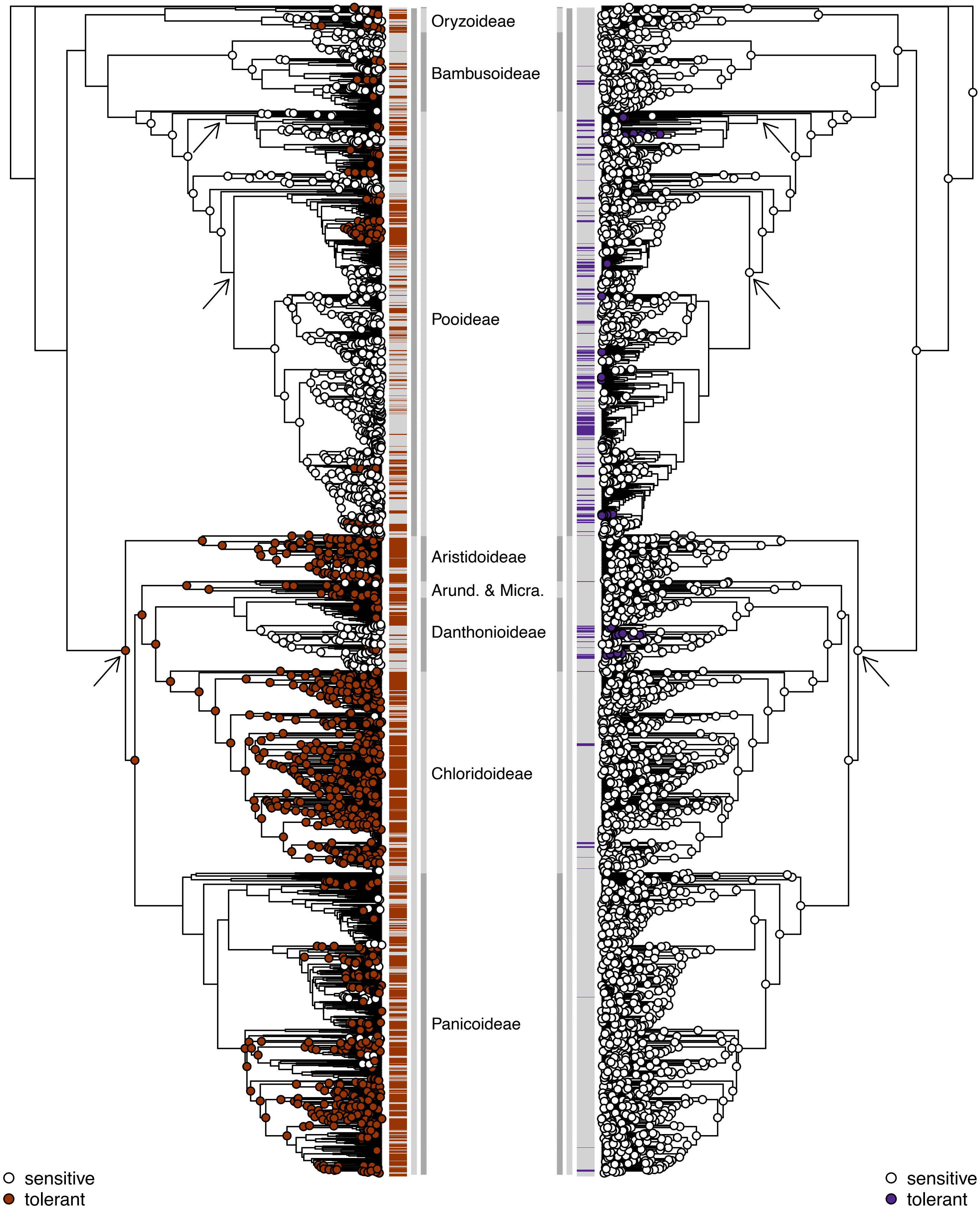

Oryzoideae
Bambusoideae
Pooideae
Aristidoideae
Arund. & Micra.
Danthonioideae
Chloridoideae
Panicoideae
sensitive
tolerant
sensitive
tolerant

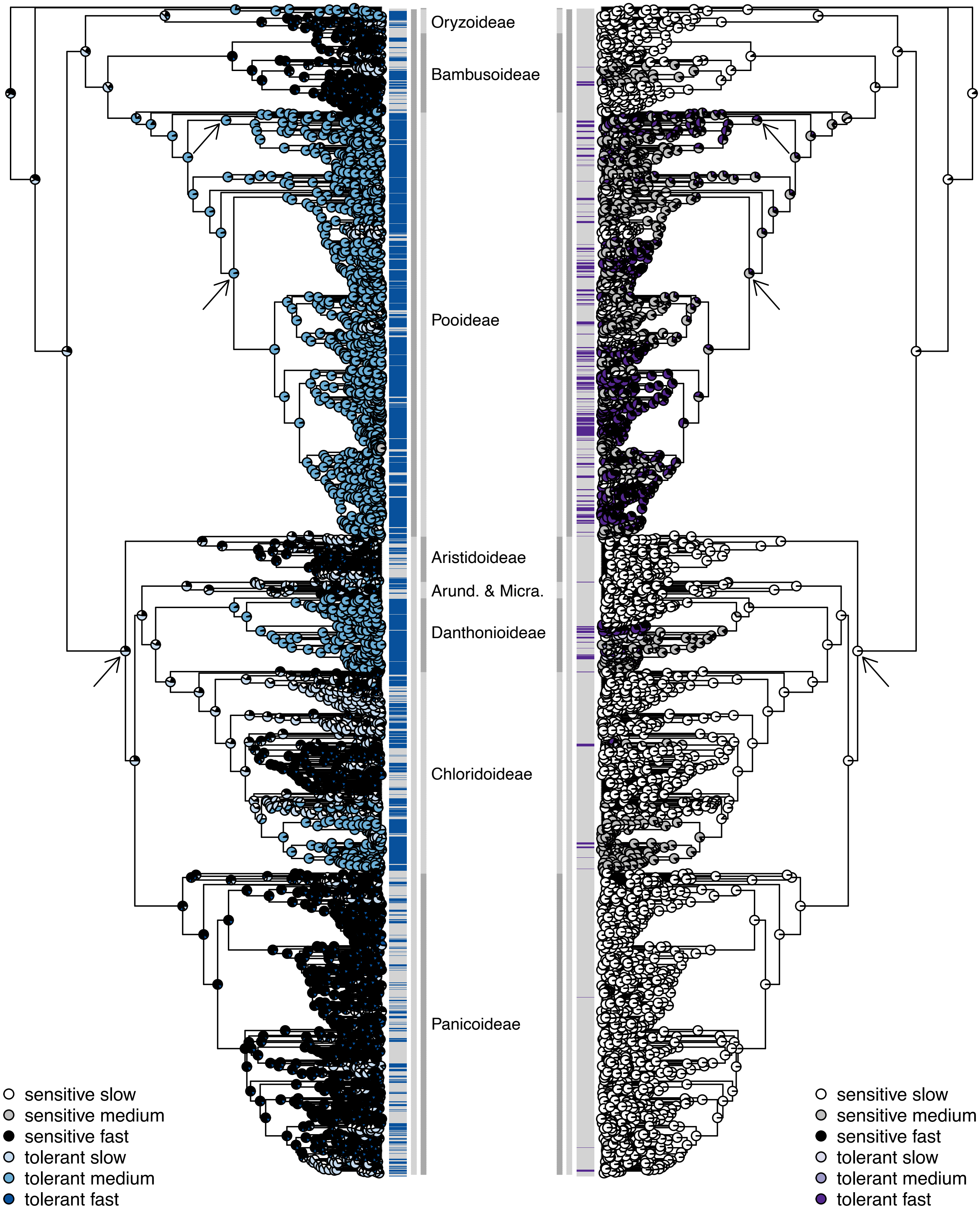
Oryzoideae
Bambusoideae
Pooideae
Aristidoideae
Arund. & Micra.
Danthonioideae
Chloridoideae
Panicoideae
sensitive slow
sensitive medium
sensitive fast
tolerant slow
tolerant medium
tolerant fast
sensitive slow
sensitive medium
sensitive fast
tolerant slow
tolerant medium
tolerant fast

**Supplementary Figure 1**. Final phylogenetic tree used in the analyses (n=2800 species sampled). Species names are shown as tip labels. Subfamilies are labelled and indicated by the arcs in greyscale. The two successive sisters to the rest (*Puelia olyriformis* and *Pharus latifolius*) are not part of any labelled subfamily. A & M = Arundinoideae and Micrairoideae.

**Supplementary Figure 2.** Schematic indicating states and transition parameters of the models used in the study. (a) Single rate independent, (b) single rate correlated, (c) single rate reduced, (d) two-rate independent and (e) two-rate reduced models. (a-c) The filled circles represent different character state combinations: white = drought and frost/winter sensitivity, red = drought tolerance and frost/winter sensitivity, blue = drought sensitivity and frost/winter tolerance, purple = drought and frost/winter tolerance. (d,e) States and transitions in the outer zone (grey) represent states and transition parameters occurring in the second, hidden rate category. Here, the filled circles also represent different character state combinations, but in a different rate category compared to the inner zone (white): black = drought and frost/winter sensitivity, dark red = drought tolerance and frost/winter sensitivity, dark blue = drought sensitivity and frost/winter tolerance, dark pink/purple = drought and frost/winter tolerance. The filled circles in the inner zone as for (a,b).

**Supplementary Figure 3.** Phylogenetic distribution of the bias in GBIF data. Tip states are coloured to represent the number of specimens per species in the tree (n=2800), defined as the number of observations following filtering and reduction to one observation per species per grid. Subfamilies are labelled and indicated by the arcs in greyscale. The two successive sisters to the rest (*Puelia olyriformis* and *Pharus latifolius*) are not part of any labelled subfamily. A & M = Arundinoideae and Micrairoideae.

**Supplementary Figure 4.** Ancestral state reconstructions for drought, frost and severe winter tolerance, displayed on facing trees for (a) drought-frost and (b) drought-winter. Ancestral states were inferred from the best-fitting hidden rates model (Tables 1,2). Ancestral state reconstructions with a marginal probability >= 0.75 are shown at each node (red = drought tolerance, blue = frost tolerance, purple = winter tolerance, white = sensitivity, irrespective of rate category). The reconstruction at unlabelled nodes is equivocal (marginal probability < 0.75 for either tolerant/sensitive). Tolerance/sensitivity of the sampled species is labelled at the tips (red = drought tolerant, blue = frost tolerant, purple = winter tolerant, grey = sensitive). Outer grey vertical line: Subfamilies ("Arund. & Micra." = Arundinoideae and Micrairoideae). Inner grey vertical line: the BOP clade (dark grey; Bambusoideae, Oryzoideae, Pooideae) and the PACMAD clade (light grey; Panicoideae, Aristidoideae, Chloridoideae, Micrairoideae, Arundinoideae, Danthonioideae). The two successive sisters to the rest sampled here (*Puelia olyriformis* and *Pharus latifolius*) are not part of any labelled subfamily or clade. Arrows indicate ancestral nodes where transitions from closed to open habitats may have occurred (Bouchenak-Khelladi et al., 2010; Elliott et al., 2023; Kellogg, 2001; Zhang et al., 2022).

**Supplementary Figure 5.** Ancestral state reconstructions for frost and severe winter tolerance displayed on facing trees. Ancestral states were inferred from the best-fitting hidden rates model (Tables 1,2). Pie charts indicate the marginal probabilities of the most likely state and rate at each node (the darker the shade, the higher the transition rate).

Tolerance/sensitivity of the sampled species is labelled at the tips (blue = frost tolerant, purple = winter tolerant, grey = sensitive). Outer grey vertical line: Subfamilies ("Arund. & Micra." = Arundinoideae and Micrairoideae). Inner grey vertical line: the BOP clade (dark grey; Bambusoideae, Oryzoideae, Pooideae) and the PACMAD clade (light grey; Panicoideae, Aristidoideae, Chloridoideae, Micrairoideae, Arundinoideae, Danthonioideae). The two successive sisters to the rest sampled here (*Puelia olyriformis* and *Pharus latifolius*) are not part of any labelled subfamily or clade. Arrows indicate ancestral nodes where transitions from closed to open habitats may have occurred (Bouchenak-Khelladi et al., 2010; Elliott et al., 2023; Kellogg, 2001; Zhang et al., 2022).
